# The effects of P2Y12 loss on microglial gene expression, dynamics, and injury response in the cerebellum and cerebral cortex

**DOI:** 10.1101/2024.09.25.614526

**Authors:** Mark B. Stoessel, Rianne D. Stowell, Rebecca L. Lowery, Linh Le, Andy N. Vu, Brendan S. Whitelaw, Ania K. Majewska

## Abstract

Despite the emerging consensus that microglia are critical to physiological and pathological brain function, it is unclear how microglial roles and their underlying mechanisms differ between brain regions. Microglia throughout the brain express common markers, such as the purinergic receptor P2Y12, that delineate them from peripheral macrophages. P2Y12 is a critical sensor of injury but also contributes to the sensing of neuronal activity and remodeling of synapses, with microglial loss of P2Y12 resulting in behavioral deficits. P2Y12 has largely been studied in cortical microglia, despite the fact that a growing body of evidence suggests that microglia exhibit a high degree of regional specialization. Cerebellar microglia, in particular, exhibit transcriptional, epigenetic, and functional profiles that set them apart from their better studied cortical and hippocampal counterparts. Here, we demonstrate that P2Y12 deficiency does not alter the morphology, distribution, or dynamics of microglia in the cerebellum. In fact, loss of P2Y12 does little to disturb the distinct transcriptomic profiles of cortical and cerebellar microglia. However, unlike in cortex, P2Y12 is not required for a full microglial response to focal injury, suggesting that cerebellar and cortical microglia use different cues to respond to injury. Finally, we show that P2Y12 deficiency impairs cerebellar learning in a delay eyeblink conditioning task, a common test of cerebellar plasticity and circuit function. Our findings suggest not only region-specific roles of microglial P2Y12 signaling in the focal injury response, but also indicate a conserved role for P2Y12 in microglial modulation of plasticity across regions.

## Introduction

Microglia, the innate immune cells of the central nervous system (CNS), actively shape neural development and maintenance. While microglia were historically studied for their roles in pathology, recent research has revealed that in the healthy brain microglia are not quiet sentinels waiting for activation, but rather active surveyors and patrollers of the CNS parenchyma ^1,2^. Under healthy or physiological conditions microglia are highly ramified cells and their processes constantly move through their local milieu, contacting neural and glial elements ^1–3^. These contacts have been shown to influence neuronal plasticity, including through the elimination of synapses and the formation of dendritic spines ^4–6^. Microglial participation in neuronal plasticity is dependent upon several chemotactic and neuromodulatory signals ^7–12^, among which one of the best described is adenosine diphosphate (ADP) sensed through the microglial P2Y12 purinergic receptor ^13–16^. P2Y12 is a G_i/o_ coupled receptor highly enriched in microglia and in the CNS, but also expressed in platelets and some peripheral tissue resident macrophages ^17–20^. P2Y12 is a vital element of the microglial sensome, and is expressed in most homeostatic microglial populations, allowing microglia to respond to extracellular ADP, a product of the hydrolysis of ATP which is released from various sources within the CNS. This sensing capacity has been implicated in many homeostatic processes, such as sensing neuronal activity ^16,21^, promoting plasticity ^13,22^, and altering behavioral responses at the animal level^22,23^. Furthermore, P2Y12 is a key sensor for injury, as extracellular ATP release can also signal a pathological event. Indeed, cortical microglia require P2Y12 to complete their chemotactic response to a focal laser injury ^13,14^. However, the question remains whether the physiological and pathological roles of P2Y12 are conserved across brain regions.

Much of our knowledge of microglia-neuron interactions and the function of microglial dynamics stems from the study of cortical and hippocampal microglia, which were originally thought to be representative of most, if not all, microglial populations. However, recent research indicates that microglia exhibit high degrees of regional heterogeneity, both in terms of gene expression and dynamic behavior ^24–27^. On a transcriptional level, cerebellar microglia express higher levels of immune related genes^25^ and exhibit a different epigenetic landscape which allows for increased phagocytosis and clearance of debris, all in the absence of classical microglial reactivity ^26^. It is hypothesized that these transcriptomic differences underly differences in the morphology and dynamics observed between cerebellar and cortical microglia. Compared to cortical and striatal microglia, cerebellar microglia are less ramified, and less dense within the cerebellar parenchyma ^24,26^. As a consequence of decreased ramification and density, cerebellar microglial processes are able to survey less of the surrounding tissue. However, the somas of cerebellar microglia translocate under homeostatic conditions, a phenomenon not yet observed in microglia of any other brain region ^24^, which may partially compensate for the reduced surveillance capacity of their processes. In the context of pathology, both cerebellar and cortical microglia can respond rapidly to focal injury, by extending their processes toward the lesion, isolating it from surrounding tissue. While both cerebellar and cortical microglia complete this response on a similar time scale, it is worth noting that due to their decreased density within the tissue, cerebellar microglial processes often travel a greater distance to the lesion center in that time frame^24^. When a more systemic inflammatory stimuli is applied, such as peripheral lipopolysaccharide (LPS), cerebellar microglia upregulate signature microglial activation genes such as *Arg1* and *Nos2* at similar levels to cortical microglia. However, ∼18% percent of upregulated genes are unique to cerebellar microglia, indicating that facets of the microglial inflammation and damage response pathways may be distinct in these cells^25^. These findings suggest that cerebellar and cortical microglia show many common phenotypes and functions but can also show distinct behaviors in both physiological and pathological conditions, which need further investigation.

To understand whether P2Y12 has similar functions in cerebellar and cortical microglia, we investigated cerebellar microglial gene expression, surveillance, injury response, and cerebellar mediated behaviors in P2Y12 knock out (KO) mice compared to wild type mice. We first determine that loss of P2Y12 affects microglial density but not distribution in the cortex and cerebellum of female but not male mice. While cerebellar microglia show distinct gene expression patterns when compared to cortical microglia, these differences are only mildly perturbed by loss of P2Y12. Using *in vivo* two-photon microscopy, we show that cerebellar microglial process and baseline somal motility, as well as the response to a local injury remain largely unaffected by the loss of P2Y12. This latter is surprising, because P2Y12 is required for a full injury response in cortical microglia. Lastly, we investigated the potential role of microglial P2Y12 in mediating cerebellar plasticity using a delay eyeblink condition paradigm, showing that loss of P2Y12 dampens the learning response. Altogether, our results suggest that some functions of P2Y12, such as its contributions to synaptic plasticity, are conserved between the cerebral cortex and cerebellum, while others, such as its contributions to injury responses, may be differentially expressed by microglia in the two brain areas.

## Results

### Loss of P2Y12 increases microglial density in the cortex and cerebellum in a sex specific manner

Using immunohistochemistry, we showed that both cerebellar and cortical microglia in CX3CR1-GFP mice express P2Y12. P2Y12 immunoreactivity is prominent throughout the membrane of GFP and Iba1-positive microglia in both brain areas and is otherwise absent (Figure 1A), confirming previous findings that P2Y12 is expressed specifically in Iba1-positive cells throughout the brain ^14,28^. Importantly, P2Y12KO CX3CR1-GFP mice, did not show P2Y12 immunoreactivity in the cortex or cerebellum, validating P2Y12 loss in the KO in both brain areas. In both genotypes microglial density appeared lower in the cerebellum than in the cortex, with cerebellar microglia being less ramified than cortical microglia (Figure 1B and 1F) in agreement with previous studies ^24^. To determine how P2Y12 loss affects microglial density and distribution in the cerebellum as compared to cortex, we analyzed microglia throughout the cortical layers of primary visual cortex (Figure 1C), and in the molecular layer (ML) and granule cell layer (GCL) of cerebellar lobule IV/V (Figure 1G). We found that loss of P2Y12 increased microglial density in the cortex and cerebellum (in both the GCL and ML), specifically in females (Figure 1C and 1G). Microglial distribution in the cortex and in both layers of the cerebellum followed a similar pattern where female P2Y12KO mice had decreased spacing of cortical microglia as measured by the nearest neighbor index (NN), commensurate with an increase in density, while males did not (Figure 1D, H). These differences in NN were likely due to the effects of P2Y12 on microglial density, as the spacing index, which accounts for microglial density ^29,30^, was not significantly different in either brain region or layer (Figure 1E, I). To determine how the loss of P2Y12 affects microglial ramification, we compared the morphology of GFP positive microglia using Sholl analysis in all layers of visual cortex and both the GCL and ML of the cerebellum (Figure 1J). Consistent with previous findings ^24–26,31^, we showed that cerebellar microglia were significantly less ramified than cortical microglia, with a maximum ∼5 intersections in the cerebellum and ∼15 intersections in the cortex. Loss of P2Y12 did not affect microglial complexity in either region in either sex (Figure 1K-L). These results indicate a minimal role for P2Y12 in determining cerebellar, as well as cortical, microglial ramification, with a sex specific role in determining microglia density both in cortex and within the layers of the cerebellum.

**Figure 1.**
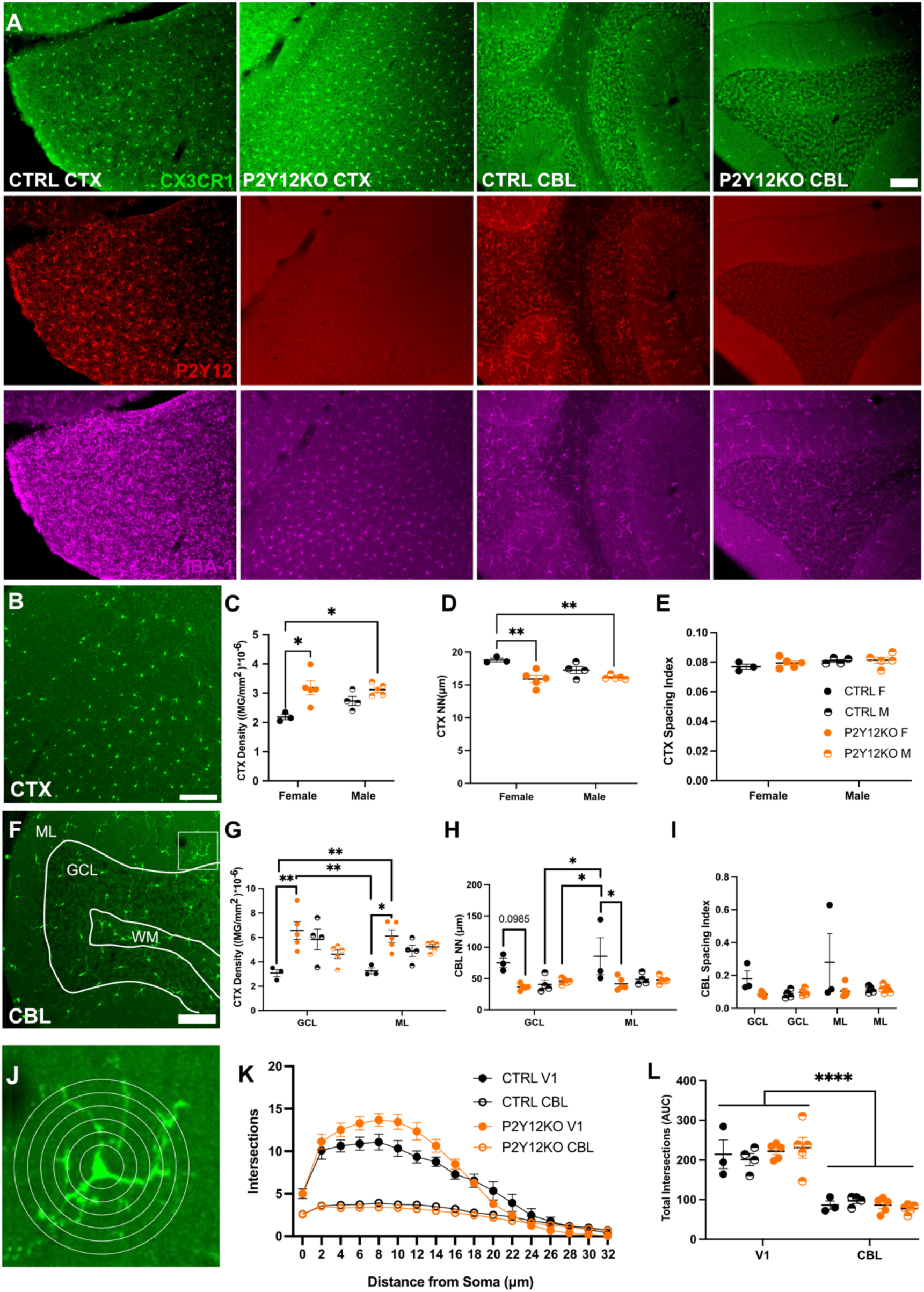
Microglial density is affected by the absence of P2Y12 in female mice. Representative confocal images of sections of visual cortex (CTX) and cerebellum (CBL), from P2Y12 deficient (P2Y12KO n=10; 5M, 5F) or control (CTRL n=7; 4M, 3F) mice, with microglial reporter CX3CR1-GFP (green) or immunohistochemical staining for microglial markers P2Y12 (red) and IBA-1 (magenta). Images confirm P2Y12 deficiency in P2Y12KO animals (A, scale bar = 100μm). Representative confocal image from CTRL CTX (B, scale bar = 50μm). P2Y12KO females but not males showed an increase in microglial density compared to CTRL (C, two-way ANOVA, significant effect of genotype, p=0.0016, interaction of sex and genotype, p =0.0985; Bonferroni post-hoc * = p<0.05). Microglial distribution, measured with a nearest neighbor index (NN) was decreased in P2Y12KO females but not in males (D, two-way ANOVA, significant effect of genotype, p=0.0006, interaction of sex and genotype, p=0.0669, Bonferroni post-hoc ** = p<0.01). Microglial spacing index showed no effect of genotype or sex in CTX (E, two-way ANOVA). Representative confocal image from CTRL CBL with boundaries between the cerebellar molecular layer (ML), granule cell layer (GCL), and white matter (WM) shown. (F, scale bar = 50μm). Boxed area is enlarged in J. GCL and ML microglial density was increased in P2Y12KO females but not in males (G, three-way matched (layer) ANOVA, significant effect of genotype, p=0.0202, interaction of sex and genotype, p= 0.0040, and interaction of layer, sex, and genotype, p=0.0044; Bonferroni post-hoc ** p<0.01). GCL and ML microglial NN indices were decreased in P2Y12KO females (H, three-way matched (layer) ANOVA, significant effects of sex, p=0.0401, genotype, p=0.0072 and interaction of sex and genotype, p=0.0039; Bonferroni post-hoc * p<0.05). CBL microglial spacing index showed no effect of genotype, sex, or layer (I, three-way matched (layer) ANOVA). Representative confocal cerebellar microglia from inset in (F) with concentric rings originating at the cell soma demonstrating Sholl analysis as a measure of microglial arbor complexity (J). The number of intersections with Sholl rings was quantified and plotted as a function of distance from soma center in each region and genotype (K). CBL microglia were significantly less ramified than CTX microglia, but ramification was not affected by sex or genotype (L, three-way matched (region) ANOVA, significant effect of region, **** p<0.0001)

### Cortical and cerebellar microglia have different protein signatures which differ between sexes and are affected by loss of P2Y12

Given the sex-specific effects of loss of P2Y12 on microglial density in both brain regions, we wanted to better understand how cortical and cerebellar microglia phenotypes change in P2Y12KO mice of both sexes. Since P2Y12KO microglia have not previously been characterized at the transcriptomic level, we chose to isolate microglia from both genotypes, sexes and brain regions using fluorescence activated cell sorting (FACS; Supplemental Figure 1) and then perform RNA sequencing analysis. Because of the sex-specific effects of P2Y12 loss on microglial density, and the technical limitations on processing the large set of samples, we performed the analysis separately in the two sexes. Microglia were isolated from the cortex and cerebellum based on CD11b+ and CD45lo expression (Supplemental Figure 1A) ^32^ and, in addition, were stained with a panel of common microglial makers for flow cytometry (Figure 2 and Supplemental Figure 1A) ^19,33,34^. P2Y12KO microglia showed a robust loss of P2Y12 in both sexes, confirming the loss of P2Y12 protein expression we reported using immunohistochemistry (Figure 1, 2A-B). Further, there were differences in P2Y12 expression between cortical and cerebellar microglia in control (CTRL) mice that differed by sex, with lower P2Y12 levels in cerebellar microglia of male mice, but higher expression in cerebellar microglia of female mice as compared to cortical microglia (Figure 2A-B). Microglial CD11b levels were higher in the cerebellum than in the cortex of male mice only, however, loss of P2Y12 decreased CD11b expression in both sexes and in both brain areas (Figure 2C-D). Similarly, CD45 levels were also higher in the cerebellum of male mice only, but P2Y12 loss decreased these levels in the cerebellum of male mice and in the cortex of female mice (Figure 2E-F). CD68 and TMEM119 were more highly expressed in the cortex than in the cerebellum in males but the opposite was true for females. P2Y12 loss did not affect CD68 or TMEM119 levels in either brain region in either sex (Figure 2 G-J). Broadly, these results align with literature descriptions of cerebellar microglial as more “immunovigilant” or macrophage like, though with some key sex-specific aspects ^25,26^.

**Figure 2.**
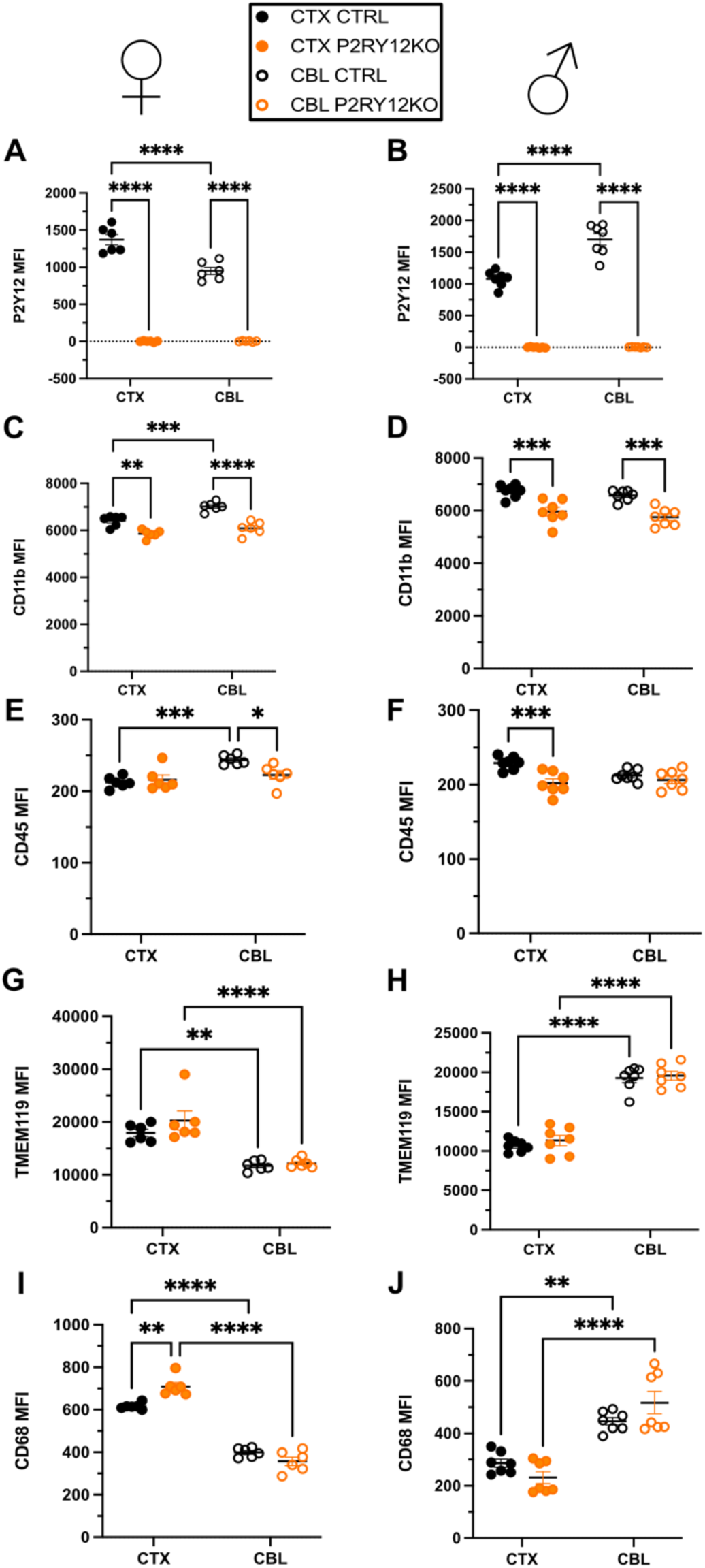
Microglial protein expression patterns are primarily controlled by region and genetic sex with some minor effects of P2Y12 deficiency. Flow cytometry of FACS sorted microglia, with common microglial makers. Microglia were sorted from the cortex (CTX, closed circles) or the cerebellum (CBL, open circles) from P2Y12 deficient (P2Y12KO, orange) or control (CTRL, black) male (left) and female (right) mice. N=7 for all groups. Results are shown as mean fluorescence intensity (MFI) for P2Y12 in male mice (two-way ANOVA, significant effects of genotype p<0.0001, region p=0.0001, with significant interaction p<0.0001; Tukey post-hoc **** p<0.0001) (A) and female mice (two-way ANOVA, significant effects of genotype p<0.0001, region p=0.0001, with significant interaction p<0.0001; Tukey post-hoc **** p<0.0001) (B); CD11b in male mice (two-way ANOVA, significant effects of genotype p<0.0001, region p=0.0002, with interaction p=0.0523; Tukey post-hoc ** p<0.01; ***p<0.0005; **** p<0.0001) (C) and female mice (two-way ANOVA, significant effect of genotype p<0.0001; Tukey post-hoc ***p<0.0005) (D); CD45 for male mice (two-way ANOVA, significant effect of region p=0.0009, with interaction p=0.0177; Tukey post-hoc * p<0.05; ***p<0.0005) (E) and female mice (two-way ANOVA, significant effect of genotype p=0.0009, with interaction p=0.0221; Tukey post-hoc ** p<0.01; ***p<0.0005) (F); TMEM119 for male mice (two-way ANOVA, significant effect of region p<0.0001; Tukey post-hoc ** p<0.01; ****p<0.0001) (G) and female mice (two-way ANOVA, significant effect of region p<0.0001; Tukey post-hoc ****p<0.0001); and CD68 in male mice (two-way ANOVA, significant effect of region p<0.0001, with interaction p=0.0002; Tukey post-hoc ** p<0.01; ****p<0.0001) (I) and female mice (two-way ANOVA, significant effect of region p<0.0001, with interaction p=0.0250) (J).

### Microglial gene expression patterns are primarily controlled by region and genetic sex with minor effects of P2Y12 deficiency

Principal component analysis (PCA) of microglial gene expression patterns obtained using RNA sequencing analysis of P2Y12KO and CTRL microglia showed limited clustering based on brain region rather than genotype in both sexes (Figure 3A and 3F). In both male and female CTRL mice, comparison of numbers of differentially expressed genes (DEGs) between regions revealed a large number of upregulated (males: 3768; Figure 3B; females: 2973; Figure 3G) and downregulated (males: 1476; Figure 3C; females: 2259; Figure 3H) genes in the cerebellum vs. the cortex. However, many of these genes overlapped with those regulated by region in P2Y12KO mice. In CTRL males, 1731 of the of the 3768 genes (46%) upregulated in the cerebellum vs. the cortex, and 521 of the 1476 downregulated genes (35%) were similarly regulated in P2Y12KO mice. In females, 38% of upregulated genes, and 51% of downregulated genes in the cerebellum vs. the cortex were common to both CTRL and P2Y12KO microglia. In fact, many fewer DEGs (<300) were found to be regulated in comparisons between CTRL and P2Y12KO microglia in either sex or brain region (Fig. 3D, E, I, J). In general, there was also less overlap between the genes that were regulated by loss of P2Y12 in the cerebellum and the cortex, although in the case of male microglia all the genes upregulated by P2Y12 loss in the cerebellum were also upregulated in the cortex where they constituted only 7% of the total upregulated genes. Together, these data suggest that P2Y12 loss has limited effect on the region-specific transcriptomic signatures of microglia, but that these limited effects may be different between cerebellum and cortex.

**Figure 3.**
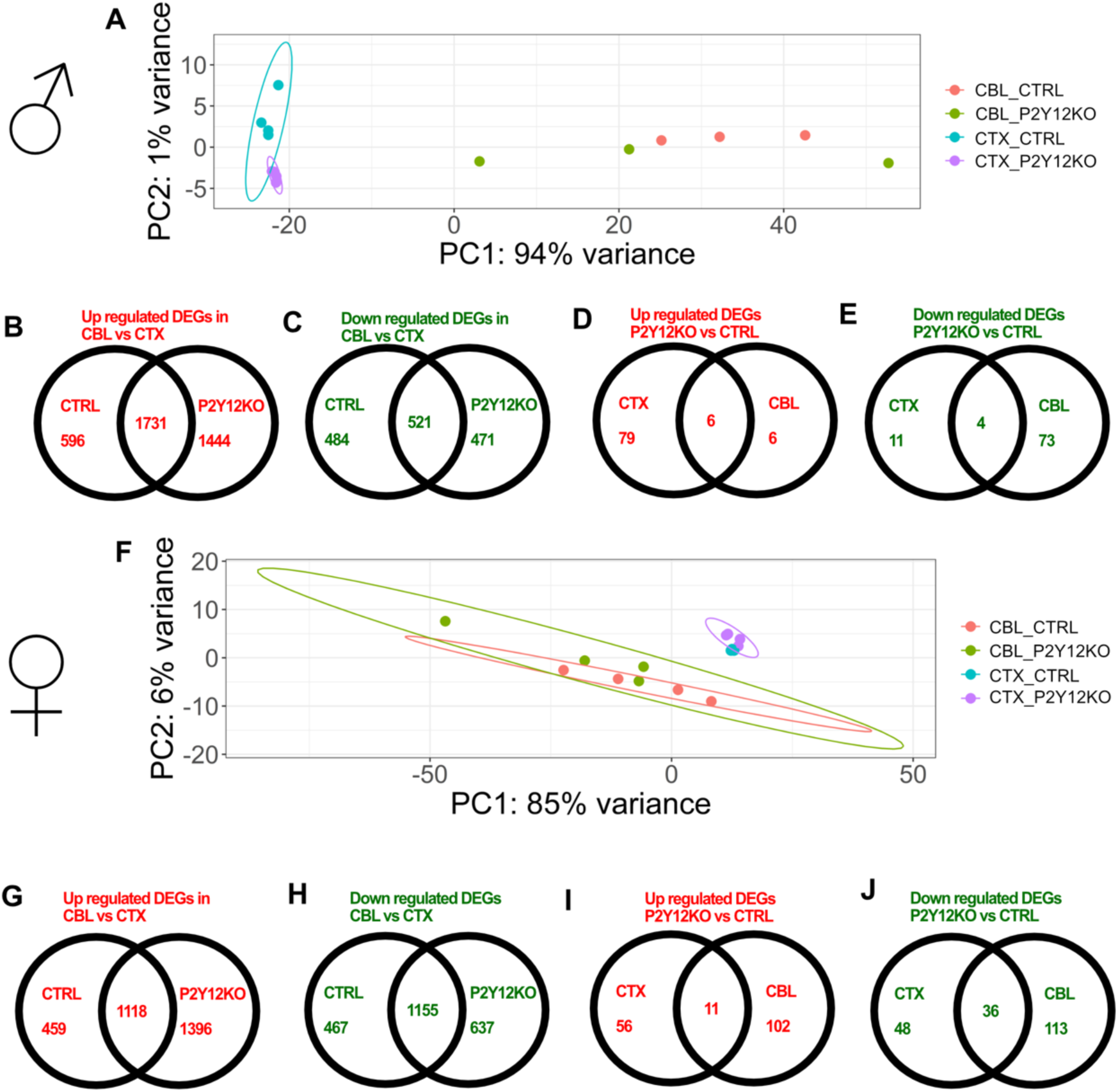
Microglial gene expression patterns are primarily determined by region and genetic sex with smaller effects of P2Y12 deficiency. Microglia were isolated from (CTRL) and P2Y12KO microglia, in both cerebellum (CBL) and cortex (CTX) and in males (A-E) and females (F-J) and profiled via RNA sequencing. Principle component analysis (PCA) of male microglia (PC1, 94% of variance, region, PC2, 1% genotype) (A). Numbers of DEGs in CBL versus CTX that were upregulated in CBL (B) and downregulated in CBL (C) in both genotypes. Numbers of DEGs in P2Y12KO versus CTRL microglia that were upregulated in P2Y12KO (D) or down regulated in P2Y12KO (E) in both brain regions. PCA of female microglia (PC1, 85% of variance, region, PC2, 6% genotype) (F). Numbers of DEGs in CBL versus CTX that were upregulated in CBL (G) and downregulated in CBL (H) in both genotypes. Number of DEGs in P2Y12KO versus CTRL that were upregulated in P2Y12KO (I) or down regulated in P2Y12KO (J) in both brain regions.

### Cerebellar microglia show increased expression of immune-related genes

To understand the sex-specific transcriptomic profiles of microglia in both brain areas, we first examined DEGs comparing CTRL cerebellar and cortical microglia in male and female mice. Broadly, our results showed microglia gene expression patterns that were consistent with the cerebellar microglial profile described in the literature ^25,26^ (Figure 4A-B, Supplementary Table 1). Specific genes upregulated in cerebellar microglia included immune response proteins *axl* in males and *axl, gvin3,* and *klf2* as well as cytoskeletal protein *myo5a* in both sexes. (Figure 4A-B). Comparisons of expression levels showed that genes that are thought to define the microglial gene signature, many of which are microglial “sensome” genes, were largely downregulated (Figure 4G and 4H Supplemental Figure 2), while immune-related genes were largely upregulated (Supplemental Figure 3) in both male and female cerebellar microglia. Gene ontology (GO) pathway^35^ analysis of DEGs showed similar changes in males and females with upregulation of genes involved in synapse organization, regulation of membrane potential, and cell junction assembly (Figure 4C, E, Supplementary Table 2) and downregulation of genes involved in autophagy, processes utilizing autophagic mechanism and leukocyte migration (Figure 4D and 4F, Supplementary Table 2). Some sex-specific differences were observed including upregulation of axonogenesis and neurotransmitter transport genes in males (Figure 4C) and RNA splicing and regulation of transmembrane ion support in females (Figure 4E).

**Figure 4.**
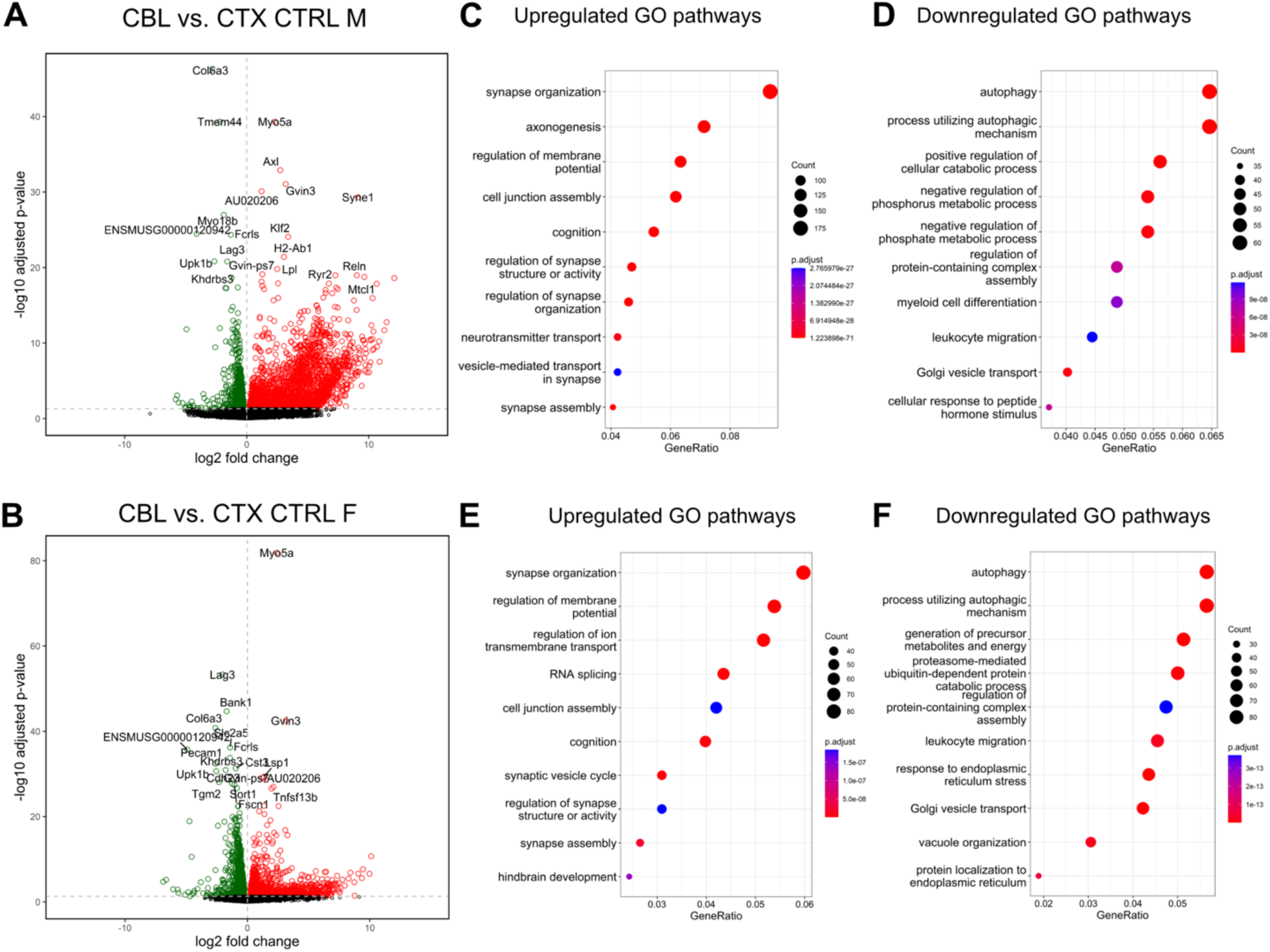
Microglial gene expression patterns differ between cerebellum and cortex in male and female control mice. Analysis of DEGs in FACS sorted microglia in the cerebellum (CBL n=3) and cerebral cortex (CTX n=4) from male and female BL6 control mice. Volcano plots of differentially expressed genes in CBL compared to CTX in males (A), and females (B) with upregulated genes in red, down regulated genes in green. Gene ontology (GO) terms upregulated (C) or down regulated (D) in male cerebellar microglia. GO terms upregulated (E) or down regulated (F) in female cerebellar microglia.

### Loss of P2Y12 does not alter the expression of microglial signature genes

Analysis of the DEGs regulated in P2Y12KO microglia revealed remarkably few differences in gene expression either in the cortex (Figure 5) or cerebellum (Figure 6), aside from the expected regulation of P2Y12 itself (Supplemental Fig. 2). Interestingly, *p2ry13* which encodes a related microglial purinergic receptor^36^ was one of the most significantly downregulated genes in P2Y12KO microglia of both sexes in both brain regions (Figure 5A-B; Figure 6A-B; Supplemental Figure 2). However, the majority of other “sensome” and microglial signature genes were not affected by loss of P2Y12 (Supplemental Figure 2). While the majority of immune-related genes were also unchanged in P2Y12KO microglia, in the cortex, Alzheimer’s Disease implicated gene *apoe* and MHC-I related transcripts were upregulated in P2Y12KO microglia of both sexes ^37,38^, while CD84 and ILI14a were downregulated (Supplemental Fig. 3). GO pathway analysis revealed that P2Y12KO cortical microglia upregulated several pathways involved in immune signaling, such as MHC signaling and antigen presentation (Figure 5C-D). Male cortical P2Y12KO microglia specifically upregulated genes involved in the defense to viruses, defense response to symbionts (Figure 5C), while female P2Y12KO microglia upregulated genes involved in the positive regulation of immune effector processes and maintenance of location (Figure 5D). Female microglia also downregulated genes involved in the regulation of the inflammatory response and negative regulator of cell activation (Figure 5E). In the cerebellum, P2Y12 loss also affected pathways that are involved in immune signaling in males (Figure 6C). However, in P2Y12KO female cerebellar microglia, where the greatest number of DEGs was observed, affected pathways did not involve immune signaling and included mRNA processing and axonogenesis (Figure 6D). In summary, our results suggest that P2Y12 loss has small effects on microglial gene expression which center around immune genes in the cortex but may affect non-immune genes in the cerebellum especially in females.

**Figure 5.**
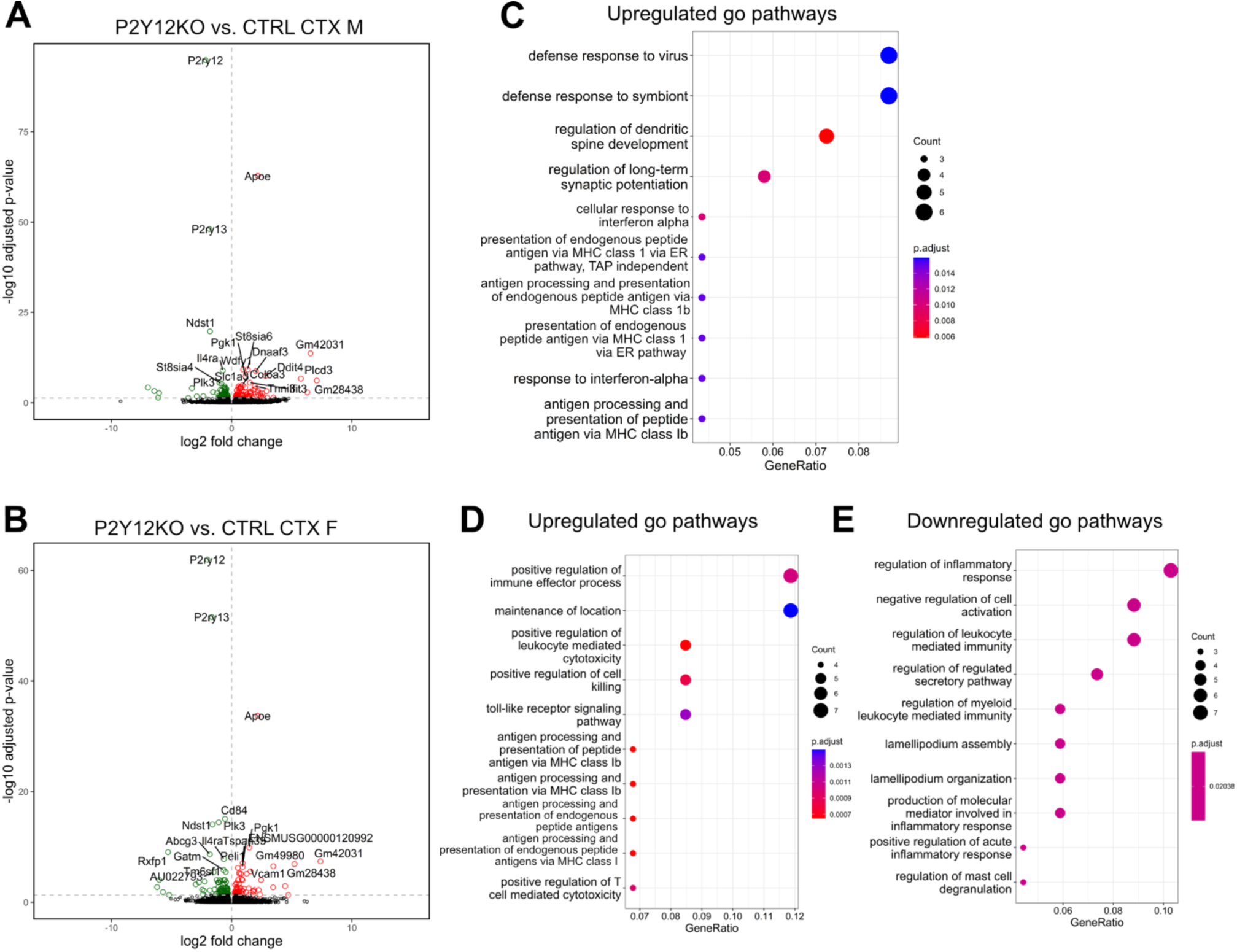
P2Y12 deficiency has subtle effects on gene expression in cortical microglia. Analysis of DEGs in FACS sorted microglia from the cortex (CTX) in male and female mice in P2Y12KO (n=4) and BL6 (CTRL n=4) mice. Volcano plots of DEGs in P2Y12KO males (A), and females (B) with upregulated genes in red, down regulated genes in green. Gene ontology (GO) terms upregulated in male P2Y12KO CTX microglia (C). GO terms upregulated (D) or down regulated (E) in female P2Y12KO CTX microglia.

**Figure 6.**
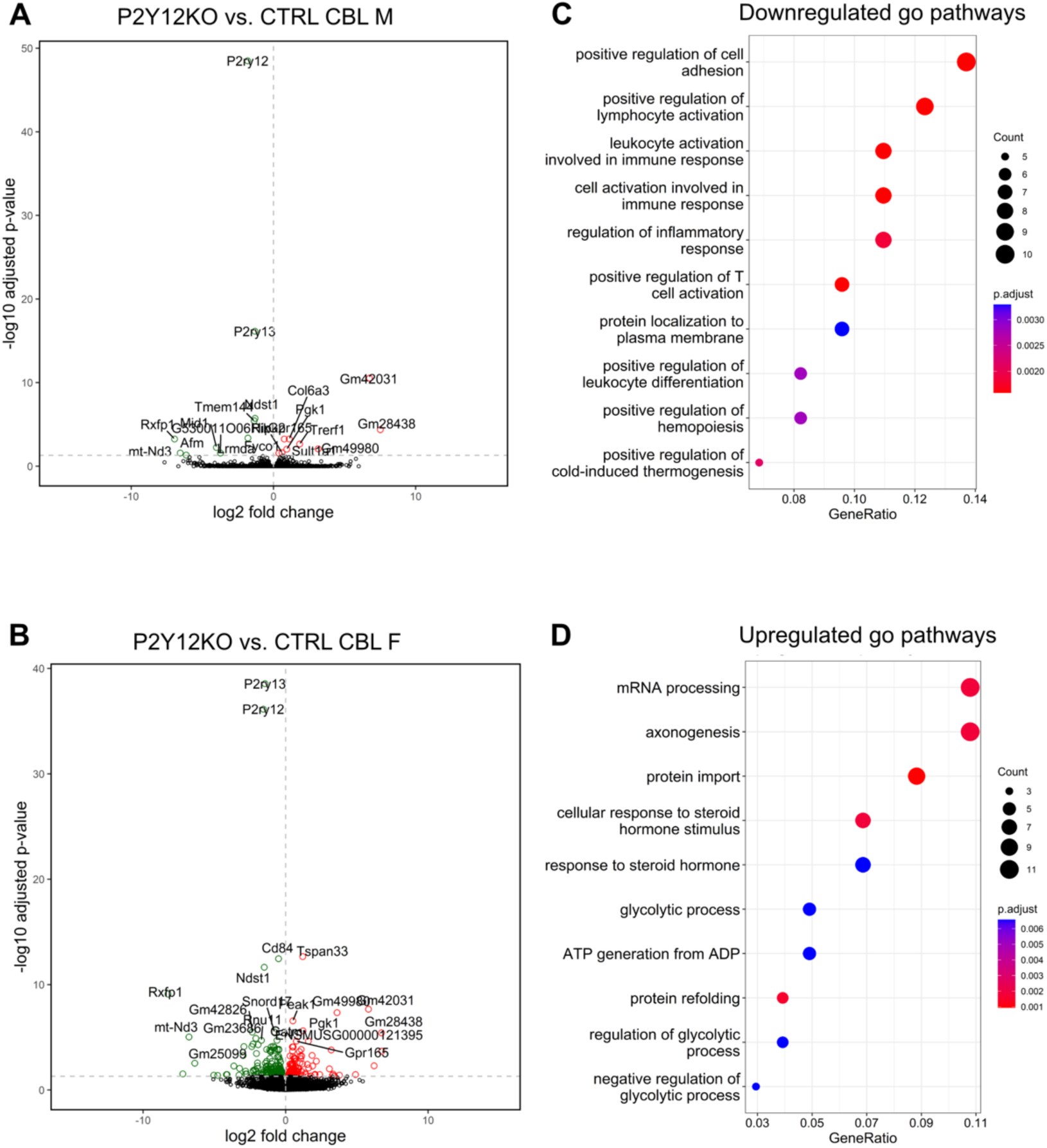
P2Y12 deficiency has subtle effects on gene expression in cerebellar microglia. Analysis of DEGs in FACS sorted microglia from the cerebellum (CBL) in male and female mice in P2Y12KO (n=3) and BL6 (CTRL n=3) mice. Volcano plots of differentially genes in P2Y12KO males (A), and females (B) with upregulated genes in red, down regulated genes in green. Gene ontology (GO) terms upregulated in male P2Y12KO CBL microglia (C). GO terms upregulated (D) in female P2Y12KO CBL microglia.

### The loss of P2Y12 does not affect the surveillance dynamics of cerebellar microglia but slightly increases cerebellar microglial somal motility

Given the effects of P2Y12 loss on the expression of immune regulators, we wondered whether P2Y12 regulates the ability of microglia to surveil the parenchyma for signs of injury and disease. To determine whether P2Y12 loss affected the dynamics of cerebellar microglia, we performed *in vivo* two-photon time-lapse imaging in anesthetized mice implanted with cranial windows over cerebellar, lobule 4/5 (Figure 7A) in either P2Y12KO CX3CR1-GFP (P2Y12KO) or CX3CR1-GFP (CTRL) controls. We then computed a microglia motility index by aligning adjacent timepoints and determining the sum of retracted and extended pixels divided by the number of stable pixels (Figure 7). This measure was then averaged for each comparison across the imaging period (Figure 7A). We found no significant differences in motility between control and P2Y12KO cerebellar microglia (Figure 7B). Similarly, the overall surveillance of microglia over the hour of imaging (obtained by superimposing images obtained at all timepoints) showed no differences between P2Y12KO and control microglia (Figure 7C). No significant differences between sexes were observed in surveillance or motility (Figure 7B-C).

**Figure 7.**
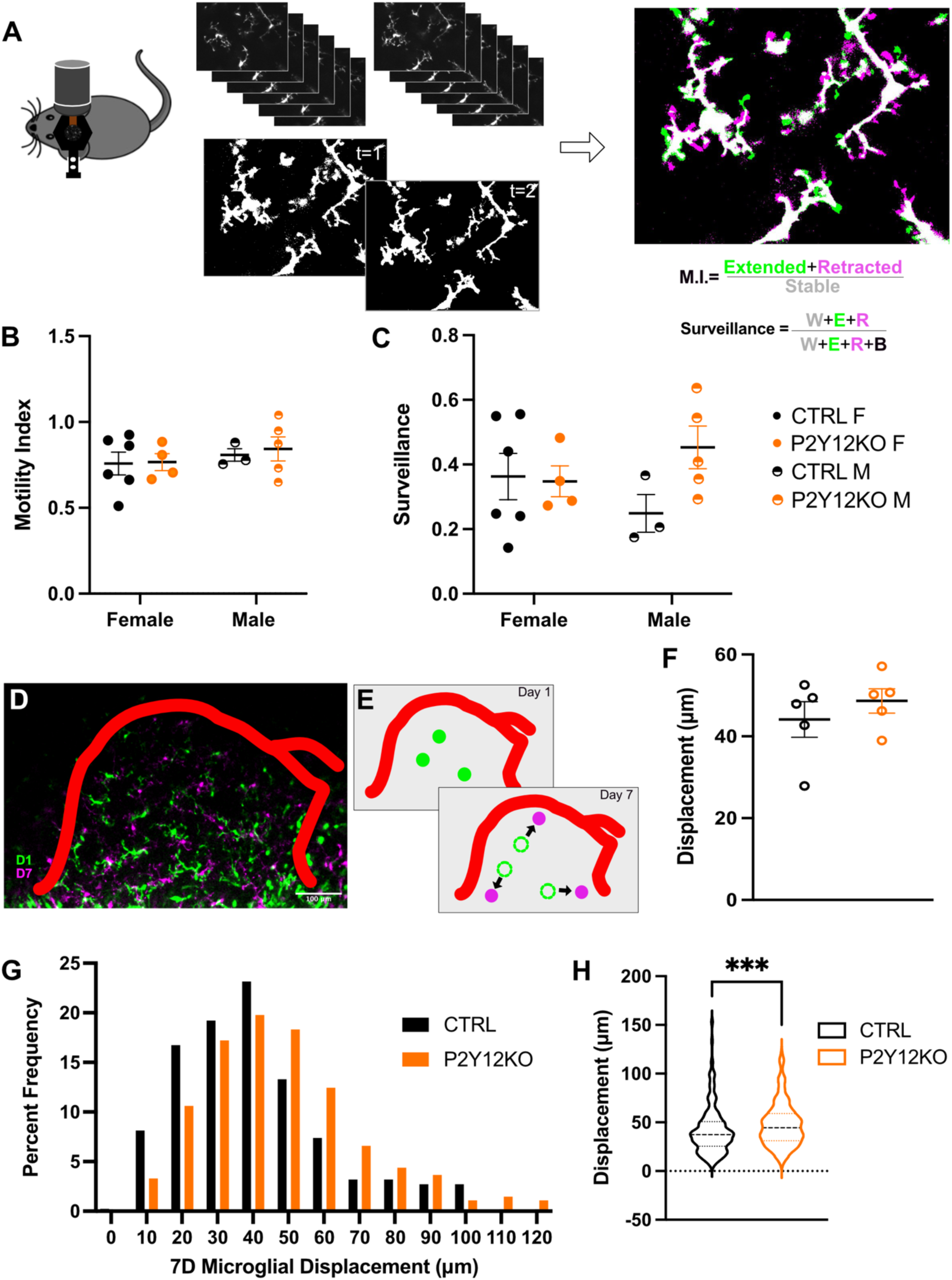
The loss of P2Y12 does not affect the surveillance dynamics of cerebellar microglia but slightly increases cerebellar microglial somal motility. P2Y12KO-CX3CR1-GFP (P2Y12KO n=9; 4F, 5M) or CX3CR1-GFP (CTRL n=9; 6F, 3M) mice outfitted with cranial windows over cerebellum (CBL) were imaged using timelapse in vivo two-photon microscopy. Mice were anesthetized then imaged once every 5 minutes for 1 hour(A). Adjacent timepoints were overlayed and extended (green), retracted (magenta), and stable (white) pixels identified (A). Microglial motility index (M.I.) was not altered in P2Y12KO mice and showed no effect of genetic sex (two-way ANOVA, Bonferroni post-hoc) (C). Similarly, microglial surveillance was not altered in P2Y12KO mice and showed no effect of genetic sex (two-way ANOVA, Bonferroni post-hoc). (C). P2Y12KO (n=5; 5F) or CX3CR1-GFP (CTRL n=5; 5F) mice outfitted with cranial windows over cerebellum (CBL) were imaged chronically at an initial day (D1, green) and one week later (D7, magenta) using in vivo two-photon imaging (D). Vasculature was used to algin images across days and soma positions were recorded. A nearest neighbor (NN) algorithm applied across the two time points allowed for measurement of putative somal displacement (E). Displacements were averaged by animal and no significant difference was found between P2Y12KO (n=5) and CTRL mice (n=5), with 50-100 microglia per animal (Welch’s t-test, two-tailed) (F). The displacements of individual microglial were plotted on a frequency distribution (G). Mean Microglial displacement (n= 273 microglia) and CTRL (n=417 microglia) was slightly but significantly increased in P2Y12KO mice (Kolmogorov-Smirnov test, p=0.0002) (H).

In addition to motile processes, cerebellar microglia have been shown to possess motile somata, in contrast with cortical microglia, whose soma remain stationary in the absence of pathology ^1,2,24,39,40^. To investigate whether P2Y12 regulates this dynamic property of cerebellar microglia, we imaged the same areas 7 days apart in mice with cranial windows and recorded somal positions on both days, utilizing vasculature patterns for alignment (Figure 7D-E). We then used a nearest neighbor algorithm between days to measure putative soma motility (Figure 7E-F see methods), a method which provides a fast way to estimate soma movement but results in values that underestimate this movement. Soma displacements over 7 days ranged from 0µm to 150µm (Figure 7G) with mean displacement of 44.12 μm in the controls and 45.30 μm in P2Y12KO, a difference which was not statistically significant (Figure 7G). When we considered the movement of each individual microglia imaged, however, we found a significant increase in the distribution of microglial displacements in P2Y12KO mice compared to controls (Figure 7G-H). All together, these data indicate that loss of microglial P2Y12 does not alter the surveillance properties of cerebellar microglial processes but may have subtle effects on cerebellar microglial somal motility.

### Cerebellar microglia respond to focal injury in the absence of P2Y12

P2Y12 is required for completion of the cortical injury response, with cortical microglia processes failing to arrive fully at the site of injury in P2Y12KO mice ^14,15^. To determine what role P2Y12 plays in the cerebellar microglia response to focal injury we created a small focal laser ablation injury *in vivo* by focusing a high-power laser beam on a single point in the imaging frame and used time-lapse imaging to capture the microglial response (Figure 8A). It is worth noting that while cerebellar microglia are less dense within the parenchyma than cortical microglia, they exhibit a robust injury response, and processes of both cortical and cerebellar microglia arrived at the ablation during the imaging period of one hour (Figure 8A-C, Supplemental Movies 1-4)^24^. To assess completeness of the ablation, we measured convergence as the microglial occupancy of the area around the ablation core at the end of the imaging session (Figures 8B-D). The convergence of control cortical and cerebellar microglia was similar, although the response in the cerebellum showed more variability than that in the cortex. P2Y12KO cortical microglia, however, failed to complete the injury response, as indicated by an average convergence of 0 (Figure 8D). The convergence of cerebellar microglia in control and P2Y12KO mice was similar in both magnitude and variability (Figure 8D) and these effects were similar in both sexes (Supplemental Figure 4A). We then measured the directional velocity of the processes, using an optical flow algorithm to measure the vector of each process between adjacent timepoints (Figure 8E-F, Supplemental Movie 5) but found no significant differences in average process velocity between groups within 10-40 minutes after ablation (Figure 8G), indicating that processes were equally motile in the direction of the injury in the presence and absence of P2Y12 in the cortex and cerebellum, despite the differences in convergence on the ablation core. This suggests that a chemotactic response is initiated in both control and P2Y12KO microglia, however, unlike cerebellar P2Y12KO microglia, cortical P2Y12KO microglia fail to reach the core. In fact, qualitative inspection of the timelapse images suggested that, in the case of P2Y12 KO microglia, cortical processes are halted well away from the core, while cerebellar microglial processes can proceed to access the injury core (Figures 8D, see Supplemental Movies 1-4). Thus, we wanted to further quantify the response not only around the core but in the whole imaging frame. We therefore measured the area bounded by the microglial processes as they form a front moving toward the core (Figure 8H). This area initially decreases in all groups, however, in P2Y12KO cortex, the area never decreases below half its initial value, consistent with an initial chemotactic response, followed by a failure to reach the core (Figure 8I). At the end of the imaging session, the size of the final area was statistically significantly different between control and P2Y12KO cortical microglia, as well as between the P2Y12KO microglia in cortex vs. cerebellum (Figure 8J). Interestingly, the final area was larger in P2Y12KO vs. control cerebellar microglia, although the effect did not reach statistical significance due to increased variability in the P2Y12KO group. Because the extent of the injury was not uniform across experiments, we assessed the final bounded area as a function of ablation core size to determine whether core size contributed to the differences in responses observed.

**Figure 8.**
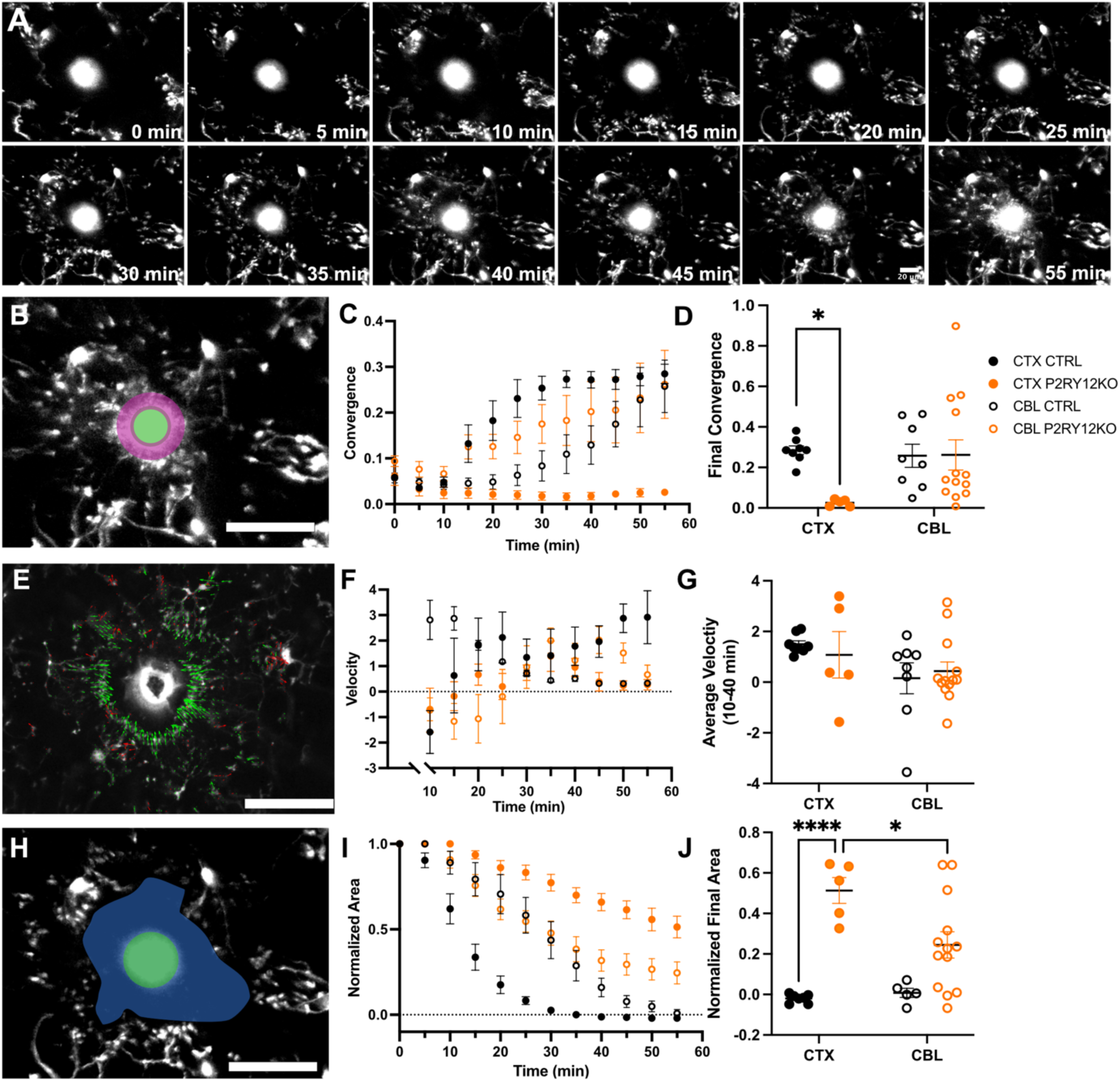
P2Y12 loss does not alter the cerebellar microglia response to focal injury. Focal laser ablation injuries were created in P2Y12KO CX3CR1-GFP (P2Y12KO) or CX3CR1-GFP (CTRL) mice outfitted with cranial windows over visual cortex (CTX), or cerebellum (CBL) and the response was monitored with time lapse two-photon microscopy every 5 minutes for 1 hour (A (CTX), scale bar = 20μm). Microglial process arrival at the injury core (green) was measured by microglial process occupancy of the ROI around injury core (magenta) (B, scale bar = 50μm). Convergence increased over time in CTRL CTX (n=8, black closed circles), CTRL CBL (n=8, black open circles), and P2Y12KO CBL (n=13, orange open circles), but not in P2Y12KO CTX (n=5, orange closed circles) (C). Convergence at the final timepoint (t=55) was lower in P2Y12KO CTX but not P2Y12KO CBL injury response (two-way ANOVA, genotype p=0.0735; interaction p=0.0648; Bonferroni post-hoc) (D). Process velocity toward (green) or away from (red) the core was measured using optic flow (E, scale bar =50μm). The velocity increased in all groups (F). Average velocity (10-40min) did not differ between groups (two-way ANOVA, effect of region 0.0605) (G). Analysis of the area bounded by processes over time (blue; H, scale bar = 50μm). Bounded area decreased in all groups over time during the microglial focal injury response (I). Area at final time point was significantly increased in the P2Y12KO CTX response (two-way ANOVA effect of region p<0.0001; genotype p=0.0824; interaction p=0.0359; Bonferroni post-hoc * p<0.05; **** p<0.0001) (J).

For all groups, except the P2Y12KO cortical group, the microglial response was not correlated with ablation core size. However, the final area in P2Y12KO cortical microglia positively correlated with ablation core size, suggesting that loss of P2Y12 resulted in chemotaxis that was sensitive to the size of the ablation (Supplemental Fig. 4B). Further, when the change in bounded area was assessed with respect to time, cerebellar microglia showed similar behavior regardless of genotype, in contrast to cortical microglia (Supplemental Figure 4C-E). Together, all our data suggests that loss of P2Y12 may alter the response of cerebellar microglia to a focal injury albeit in a much more subtle way than in cortical microglia. Thus, we argue for differing roles of P2Y12 in the cerebellar microglial injury response than in the cortical microglia injury response.

### Cerebellar microglia do not respond to acute ATP application in the absence of P2Y12

ATP has been shown to be a reliable chemotactic signal in cortical microglia only when P2Y12 is intact ^2,14,15^. The cerebellar microglial response to exogenous ATP has not been described and given that the injury response can occur in the absence of P2Y12 in the cerebellum, we hypothesized that cerebellar microglia may respond to ATP independently of P2Y12. We therefore applied exogenous ATP as a point source through a micropipette in acute cortical and cerebellar brain slices of CTRL and P2Y12KO mice and then imaged the response using two-photon time-lapse microscopy every minute for 30 minutes after application (Figure 9A-B). We established that cerebellar microglia exhibit a chemotactic response to ATP similar to that of cortical microglia and that this response is complete (i.e., the microglial processes have arrived at the ATP source, the pipette tip) by 30 minutes after application in cerebellar slices (Figure 9B), similarly to cortical microglia ^15^. To quantify the ATP response, we measured both microglial process convergence at (Figure 9C-H) and velocity toward (Figure 9I-N)^15^ the pipette tip. While convergence of microglial processes on the pipette tip was similar in cortical and cerebellar CTRL slices at the end of the imaging session, we observed a significant decrease in the convergence in P2Y12KO slices compared to control slices in both regions (Figures 9D-H), though this difference in convergence does not become significant until the later portion of the response (24 minutes post application, Figure 9H). Process velocity was also affected by P2Y12KO in both regions early in the response (6 minutes post application, Figure 9L) but not late in the response (24 minutes post application, Figure 9N) when the control response to ATP was complete. At an intermediate timepoint (15 minutes post application; Figure 9M), only cerebellar velocity was significantly different between groups (Figure 9J). This suggests that loss of P2Y12 results in a lack of initiation of chemotaxis towards an ATP point source in both brain regions, resulting in a lack of convergence onto the point source by the end of the imaging period. The response to ATP appears to be prolonged in the cerebellum likely as a result of decreased microglial density which requires microglial processes to travel a larger distance to converge on the pipette. Overall, we found that both cortical and cerebellar microglia require P2Y12 in order to respond to a point source of ATP, indicating that the differential injury response we observed in the cortex and cerebellum of P2Y12KO mice *in vivo* (Figures 8D and 8J) is not mediated by ATP signaling through another receptor in cerebellar microglia but more likely through another chemotactic pathway that is not active in cortical microglia.

**Figure 9.**
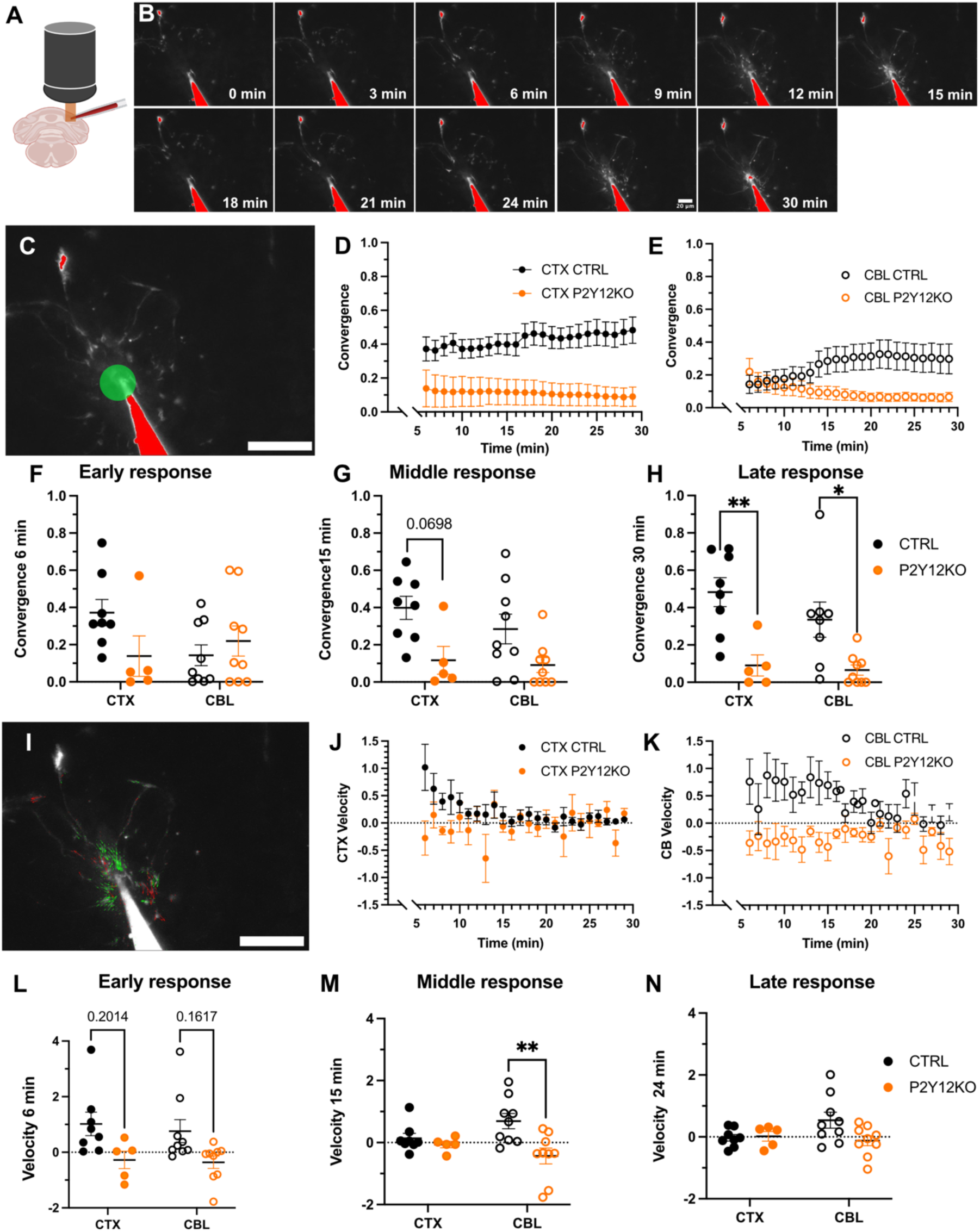
Both cerebellar and cortical microglia do not respond to acute ATP application in the absence of P2Y12. Acute slices of cortex (CTX) and cerebellum (CBL) were prepared from P2Y12KO C3XCR1-GFP (P2Y12KO CBL n=9; CTX n=5) and CX3CR1-GFP (CTRL; CBL n=9, CTX n=8) mice (A). ATP was applied to the slice through a micropipette as a point source and microglial processes converge on the pipette tip in a similar manner to the focal injury response. The microglial response was monitored using time-lapse two-photon microscopy, imaging every minute for 30 minutes (B, CBL CTRL shown, scale bar=20μm). Microglial process arrival at the pipette tip was measured by quantifying occupancy of microglial processes around pipette tip (convergence, green) (C, scale bar =50μm). Convergence increased over time in control but not P2Y12KO slices in CTX (D) and CBL (E). The early microglial response (t=6) showed no significant differences between region or genotype (two-way ANOVA, interaction p=0.0581; Bonferroni post hoc) (F). By the middle of the microglial response (t=15) there was a significant effect of genotype driven largely by the loss of the response in P2Y12KO CTX (two-way ANOVA significant effect of genotype, p=0.0015; Bonferroni post hoc) (G). The late microglial response (t=24) showed a significant effect of genotype in both CTX and CBL (two-way ANOVA, significant effect of region, p<0.0001, Bonferroni post hoc * p<0.05; **p<0.01) (H). Microglia process velocity toward the pipette tip was measured using optic flow (I, scale bar = 50μm). Microglia process velocity decreased over the course of the response in both CTX (J) and cerebellum (K) of CTRL but not P2Y12KO mice. Early process velocity (t=6) showed a significant effect of genotype (two-way ANOVA, effect of genotype p=0.0034; Bonferroni post hoc **p<0.01) (L). The middle response(t=15) also showed a significant effect of genotype, largely driven by the continued CTRL CBL response (two-way ANOVA, significant effect of genotype p=0.0079; Bonferroni post-hoc ** p<0.01) (M) At the late response time point (t=24) there were no significant effects, which indicated a completion of the response (two-way ANOVA, Bonferroni post hoc) (N).

### Cerebellar learning and plasticity are attenuated in the absence of P2Y12

Having observed differences in the cerebellar microglial injury response, we next wanted to investigate the potential role of P2Y12 in cerebellar learning and plasticity. To do so, we assessed performance on a delayed eyeblink conditioning (dEBC) task in P2Y12KO and CTRL male mice. dEBC is a well-accepted, commonly used paradigm to assess cerebellar plasticity and associative learning, and unlike trace EBC, dEBC has been shown to be dependent on the cerebellum^41–44^. Sex differences have been observed in dEBC learning rates, and male mice were chosen as the slightly slower learning kinetics of males allow for better detection of subtle differences between groups ^45^. During the dEBC task, a neutral, conditioned stimulus (CS, an LED at 405nm) is paired with a noxious, unconditioned stimulus (an air puff to the cornea at 20psi) and an association is learned between the two stimuli, such that over the course of ten days mice learn to blink in response to the CS, in anticipation of the US (Figure 10A-B). We conducted 220 such trials per day per mouse in male P2Y12KO and control mice and measured percent conditioned response (CR) to assess cerebellar learning in the presence and absence of P2Y12 ^42,46^. Control mice were able to learn the association and exhibited a percent CR of 85% by day 6. P2Y12KO mice initially seemed to learn at a rate similar to that of control mice but, by the end of the paradigm, achieved a significantly lower percent CR than controls at 65% at D10 (Figure 10C). This suggests that loss of P2Y12 attenuates cerebellar learning and plasticity in adult male mice.

**Figure 10.**
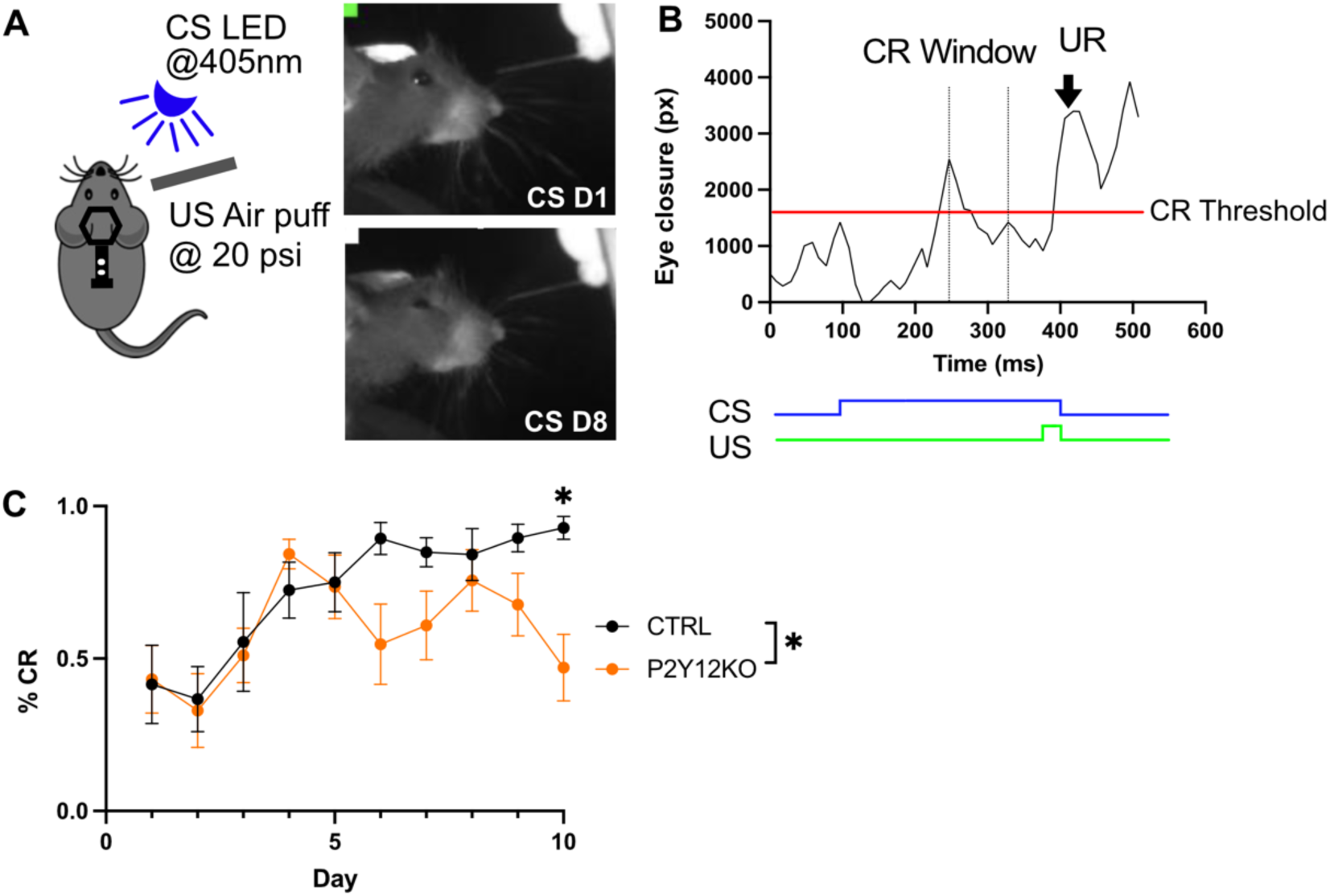
Cerebellar learning and plasticity are attenuated in the absence of P2Y12. Schematic showing dEBC conditioning paradigm pairing a neutral, conditioned stimulus (CS, an LED at 405nm) and a noxious, unconditioned stimulus (US, an air puff at 20 PSI to the cornea). On day 1 (CS D1) there is no response to the CS alone, but by day 8 (CS D8) the mouse has learned to blink in response to the CS in anticipation of the noxious US (A). Video oculography of a single successful trial plots eye closure as a function of time with CS (blue) and US (green) epochs shown below. The unconditioned response (UR) is also shown (B). Male BL6 (CTRL, black, n=7) or P2Y12KO (orange, n=7) mice underwent the conditioning, and the total percent conditioned response (%CR) was compared on each day. Learning was achieved in both groups by day 5 as both significant effects of time and genotype were found (two-way mix model ANOVA with repeated measures, significant effects of time p<0.0001; and time and genotype p=0.0355, Bonferroni post hoc * p<0.05) (C).

## Discussion

Microglia, as the primary immune cell of the CNS, exist in a continually surveillant state sampling their local environments through a variety of surface receptors and channels ^1,18^. Microglia are known to be sensitive to local neuronal damage and injury ^47^, long range neuromodulatory cues and behavioral states ^7,12^, patterns of neuronal firing ^16,40^ and behavioral experience ^3^. The P2Y12 receptor and microglial purinergic signaling are thought to comprise a key element of the microglial sensome and play a role in many of the interactions noted above ^13,16,18,19,23,28^. Here we show that the P2Y12 receptor contributes to the function of cerebellar microglia, albeit not in the exact same manner as in cortical microglia, where it has been more extensively studied. We report sex specific effects of P2Y12 deficiency on cerebellar and cortical microglia density and distribution, but no effects on microglial process arbor complexity. Transcriptomic analyses of purified microglia from cortex and cerebellum, showed large differences in the expression patterns of these two microglial populations, but a much smaller effect of P2Y12KO in either brain region which centered largely on immune signaling pathways. Dynamically, we observed few effects of P2Y12KO in microglial motility and surveillance but a subtle effect on cerebellar microglial somal motility. Loss of P2Y12 affected the responses of both cortical and cerebellar microglia to focal injury, but this effect was more profound in the cortex ^13,14^. The ability of cerebellar microglia to respond to injury in the absence of P2Y12 was not due to the sensing of ATP through a different purinergic receptor expressed in cerebellar microglia, as loss of P2Y12 abolished the response to exogenous ATP application equally in both brain regions. Finally, we demonstrated that the absence of P2Y12 attenuates cerebellar learning and plasticity implicating a behaviorally relevant role for microglia in modulating plasticity and behavior in the physiological, adult, cerebellum. Overall, our results indicate a conserved function of P2Y12 in plasticity in the cerebellum and cortex, but a differential function in its contribution to the dynamics of the focal injury response despite the fact that P2Y12 loss has only small effects on the microglial transcriptome.

### P2Y12 deficiency and regional microglial gene expression patterns

P2Y12 is an integral protein in the microglial sensome. It has been labeled a canonical homeostatic microglial marker ^14,19,28^ and is thought to be central to microglial interactions with neurons and to animal behavior ^13–15,23,40,48^. Interestingly, however, the effects of P2Y12 deficiency on the transcriptome of microglia have never been assessed. Here, we performed RNA-sequencing and flow cytometry on FACS sorted cortical and cerebellar P2Y12KO microglia from male and female mice and compared them to control mice. Given the role of P2Y12 in microglial function, it was surprising that we saw relatively few changes in overall microglial gene expression patterns in P2Y12KO microglia. While cortical and cerebellar microglia showed very different expression patterns, confirming previous literature ^25,26^, loss of P2Y12 resulted in 10-fold smaller numbers of DEGs with predominantly immune pathways affected. Loss of P2Y12 appeared to have the greatest effect in female microglia of both regions but particularly in female cerebellar microglia, which exhibited the highest number of DEGs. GO term analysis of DEGs in female cortical P2Y12KO microglia showed a down regulation of genes involved in lamellipodium assembly and lamellipodium organization, genes which are involved in assembling the leading edge of immune cell migration. Although cortical microglia soma motility is rare under homeostatic conditions and decreases with loss of P2Y12 ^40^, it is possible that these lamellipodia related genes are responsible for the differences in microglial density, soma translocation and injury response in P2Y12KO microglia, although it is puzzling why these pathways only appear in the analysis of female cortical microglia. P2Y12 loss upregulates intracellular pathways in cerebellar microglia, such as those related to mRNA and protein processing, glycolysis and axonogenesis, which may alter microglial state and induce changes in microglial responses. In summary, our results point to different transcriptional patterns in cerebellar microglia and cortical microglia, with smaller effects of P2Y12KO, which appear to affect different pathways in cerebellar and cortical microglia.

### The role of P2Y12 in maintaining cerebellar microglia morphology, density, and dynamics

P2Y12KO mice have been reported to have sex-specific increases in microglial density and altered microglial ramification ^49^. Here, we report a female specific increase in microglial density in both cerebellum and visual cortex in P2Y12KO mice but no change in microglial ramification in either cortex or cerebellum. This can perhaps be explained by the fact that our measurements are taken in two dimensions rather than three and focus on only one measure of microglial morphology. We show that cerebellar microglia are ramified and surveillant but that this process is P2Y12 independent, similar to what has been described in cortical microglia ^13,14,28^. Cerebellar microglial somas can translocate, unlike cortical microglia, and such translocation has been thought to allow the less ramified cerebellar microglia to achieve similar surveillance rates to cortical microglia ^24^. Cerebellar microglial somal motility may also be facilitated by increased expression of immune response genes by cerebellar microglia under basal conditions ^25,26^, although the mechanisms by which immune signaling contributes to cerebellar microglia soma motility remain poorly understood. Our results indicate a subtle increase in somal motility in the absence of P2Y12, suggesting that purinergic signaling, which is heavily implicated in directed microglial chemotaxis and in microglial process interactions with neurons ^2,14–16,21,48^, limits microglial soma movement. It is interesting to note that the more subtle somal translocations of cortical microglia, were described to decrease in the absence of P2Y12, and the increased somal motility of hippocampal microglia during epileptiform activity is also dependent on P2Y12 signaling ^40^. Thus, P2Y12 is implicated in somal translocation in cortical, hippocampal, and cerebellar microglia in health and disease, albeit in opposite ways. This suggests that the role of P2Y12 in microglial soma translocation merits further, more rigorous study and suggests that control of microglial somal motility may proceed by different mechanisms in different brain areas. Microglia may therefore rely upon localized ATP released during neuronal and perhaps glial activity for guidance during somal translocation, though transduction of this signal between cortex and cerebellum leads to differing overall downstream effects on somal motility. If so, microglial somal movement throughout the cerebellum is likely much less random than previously thought ^24^. P2Y12 might be necessary as a stop signal to ensure that microglia stay at sites of neuronal activity, or alternatively, the loss of P2Y12 could push cerebellar microglia toward a more motile state, increasing their random movements. While the effects of P2Y12 loss on the transcriptome of cerebellar microglia were subtle, upregulation of cytoskeletal genes such as *myo5a* in P2Y12KO microglia could be involved. Though poorly understood in microglia, *myo5a* has been shown to be involved in non-muscle cell movement in other cell types ^50^. Thus, understanding the role of *myo5a* and other cytoskeletal proteins in cerebellar homeostatic somal motility could be a fruitful area of study ^51,52^.

### P2Y12 and microglial chemotaxis to focal injury

One of the best described pathways of chemotaxis of cortical microglia processes is dependent on P2Y12 and involves microglia reorientating their arbors to rapidly move their processes toward sites of focal ATP release ^2,14,15,53^. In the absence of P2Y12, cortical microglia appear to initiate chemotaxis but never complete their response, instead pausing or getting “stuck” at a significant distance from the injury. This distance scales with the size of the ablation which suggests that some aspect of the ablation itself acts as a stop signal arresting microglial chemotaxis only in the absence of P2Y12. EM images of the ablation core show that damage to the brain tissue occurs outside of the ablation core (Supplementary Fig. 4F-G) and P2Y12 may be required for cortical microglia to proceed past this radius of damage to reach the core.

The fact that an initial chemotaxis occurs in cortical microglia in the absence of P2Y12 suggests that these microglia may be sensitive to other cues released in the injury process. In fact, microglial process chemotaxis in the cerebellum is largely P2Y12 independent. While loss of P2Y12 did affect the movement of the front of cerebellar microglial processes towards the core, the effect was subtle and other measures of microglial response, such as convergence and process directed velocity were largely normal in P2Y12KO cerebellum. The mechanisms underlying the P2Y12 independent response of cerebellar microglial are likely highly complex and multivariate and may only partially overlap with those in the cortex. One possibility is that the baseline transcriptional state of cerebellar microglia, which generally involves heightened expression of immune mediators as described above, allows for increased involvement of alternative injury sensing pathways in eliciting directed microglial chemotaxis. The cerebellar microglial transcriptional program supports increased phagocytosis and thus, it is possible, supports differential injury response mechanisms than those seen in cortical microglia ^26,54^. It should be noted that microglia express other purinergic receptors, such as P2Y13 and P2X7 ^36,55^. While the simplest explanation is that P2Y12KO cerebellar microglia can respond to ATP gradients utilizing one of these alternate purinergic signaling pathways, our ATP application experiments suggest this is not the case, as ATP did not elicit the chemotactic response seen in P2Y12KO cerebellar microglial during the focal injury response. Given the speed and scale of the cerebellar microglial injury response, we hypothesize that the factors guiding this response in the absence of P2Y12 must be, like ATP, rapidly diffusible, released during and immediately after the laser ablation, and elicit a response in P2Y12KO and control cerebellar microglia. What these factors are and their specificity to the cerebellum remain exciting avenues for future study.

### The contributions of P2Y12 signaling to learning and synaptic plasticity

It is known that microglia participate in neuronal plasticity in many brain regions, although such participation is often limited to developmentally relevant critical periods. For example, microglia have been shown to participate in shifts in ocular dominance in visual cortex ^7,13^ (although the role of microglia in this process remains unclear^56^), thalamocortical input segregation in barrel cortex ^9,10^, and retinogeniculate input segregation ^4,5,57^, all through a variety of molecular mechanisms and signaling pathways. Hippocampal microglia have been shown to modulate neuronal long-term potentiation (LTP) and long-term depression (LTD) well into adulthood through cytokine secretion or synaptic phagocytosis ^58–60^. Cerebellar microglial modulation of cerebellar circuits has been described, but in the context of neuroinflammation and microglial activation ^61^. The cerebellum is a highly plastic brain region that continually adapts to changing behavioral circumstances ^62^. At the synaptic level, this manifests in a delicate balance of LTP and LTD at the cerebellar climbing fiber-Purkinje cell synapse and the parallel fiber-Purkinje cell synapse. These two synapse types are the key excitatory inputs to the cerebellar Purkinje cell, which is itself the primary output cell of the cerebellum ^62^. Deficits in either LTP or LTD in the cerebellum have been shown to disrupt cerebellum function and learning ^63–65^. Here we show that mouse performance in a dEBC paradigm is attenuated in the absence of P2Y12, suggesting a role for microglia in modulating cerebellar plasticity and learning in the adult brain (Figure 10).

In other brain regions, purinergic and adenosine signaling have been implicated in both microglia participation in synaptic plasticity and in microglial modulation of neuronal activity. As mentioned above, P2Y12 signaling contributes to shifts in ocular dominance plasticity (ODP), and while shifts in ODP are highly developmentally regulated ^13^, the loss of P2Y12 can alter behavior at the whole animal level in adulthood ^22,23^. It is possible that cerebellar microglia respond to Purkinje cell firing, and their processes are recruited by ATP release from the Purkinje cell somata and dendrites, as has been described in the cortex and striatum, with these contacts modulating neuronal activity ^16,21^. Indeed, microglia in the adult cerebellum have been shown to dynamically interact with and come into close physical contact with both Purkinje cell somata and dendrites *in vivo*, though the function of these contacts is not known ^24^. It is also worth noting that many areas of the cerebellum have been implicated in dEBC, notably lobule IV/V, which we image here, but also the deep cerebellar interpositous nucleus, with inactivation or ablation of these areas causing varying deficits in dEBC learning ^66–68^. As such, there is the possibility that microglia interact with the cerebellar circuit at the level of the deep cerebellar nuclei (DCN). Very little is known about the microglia in this region, and this would be an interesting area to explore in the future. Cerebellar learning during dEBC has been shown to be dependent upon perineuronal nets (PNNs) within the DCN; when their degradation is prevented, learning is inhibited, but when it is facilitated, learning is accelerated ^69^. Intriguingly, microglia in the cortex have been implicated in the degradation of PNNs during neuronal plasticity, where clopidogrel, a potent P2Y12 inhibitor, blocks removal of PNNs ^70^. Therefore, taken together, we propose that microglia may be acting either at the level of Purkinje cell to modulate circuit activity or at the level of the deep cerebellar nuclei by affecting PNN remodeling. These mechanisms are not mutually exclusive but do present intriguing future avenues of study in discerning the role of cerebellar microglia in cerebellar behavior and plasticity at different levels of the cerebellar circuit.

In conclusion, our results suggest that microglia may tune their reliance on purinergic signaling depending on the context. Our results highlight the incredible adaptability of microglia and their ability to embed themselves in milieus that differ in cytoarchitecture, neuronal firing patterns, and behavioral roles, and further emphasize the importance of studying these cells in their context.

## Materials and Methods

### Animals

All experimental protocols were performed in strict accordance with both University of Rochester committee on Animal Resources and National Institutes of Health Guidelines. Adult mice, postnatal day (P) 60-100, were bred on a C57/BL6J background in house. A constitutive P2Y12 knockout (P2Y12KO) mouse ^13^ lacking both copies of the *p2ry12* gene was used during delay eye blink conditioning assays and was bred with CX3CR1-GFP animals (JAX 005582) ^71^ for microglial visualization during in vivo two-photon imaging experiments, immunohistochemical experiments, and in in fixed tissue analyses. Unless otherwise noted, animals were housed in 12hr light/12hr dark cycle with chow ad libitum. Experiments were performed on both male and female animals, with sex differences noted and displayed where found.

### Fixed tissue preparation

For analysis of cerebellar microglial density, morphology, and distribution both P2Y12KO CX3CR1-GFP/+ and CX3CR1-GFP/+ control animals were euthanized with an overdose of sodium pentobarbital and transcardially perfused with 0.1 M phosphate buffered saline (PBS) followed by 4% paraformaldehyde (PFA). Brains were extracted then allowed to post fix overnight in 4% PFA. Brains were then cryoprotected in 0.2M PBS 30% sucrose solution. Sagittal sections of 50μm thickness were cut on a freezing microtome, then stored in cryoprotectant (25% 0.2 PB, 25% glycerol, 30% ethylene glycol, 20% ddH2O) at 4°C. Sections were selected, briefly washed with 0.1M phosphate buffer (PB), mounted on slides (Lecia), and coverslipped with Prolong Gold mounting media with DAPI Molecular Probes, P36935).

### Immunohistochemistry

To confirm loss of P2Y12 protein, select P2Y12KO CX3CR1-GFP/+ sections underwent immunostaining for P2Y12 and IBA-1. Briefly, cortical, and cerebellar and sections were rinsed in room temperature (RT) 0.1 M PBS (three 10-min washes) and blocked in 10% bovine serum albumen (BSA) and 0.4% triton. Sections were then incubated in a primary antibody solution consisting of 0.5% BSA and 0.4% in 0.1M PBS, overnight (minimum 16h) at 4°C on a shaker in a humidified chamber (goat anti-IBA-1 1:1000, rabbit-anti-P2RY12 1:2000,). The sections were then washed with 0.1 M PBS and incubated 4h at RT in secondary antibody (Alexa-Fluor 594 1:500, donkey anti-rabbit; Invitrogen A21207). Sections were mounted onto glass slides with coverslips using Prolong Gold (Molecular Probes, P36934).

### Microglial arbor complexity

Sagittal sections containing layers 1-6 of primary visual cortex or the molecular and granular cell layers (ML and GCL, respectively) of the cerebellum were imaged using an Inverted Leica DMi8 Microscope and captured with LAX Software. Sections were stained as described above. A 20x/0.75 apochromat magnification objective, and 1024 x 1024-pixel z-stacks with a 1μm step size were collected. Images of the cerebellum and cortex were collected under a 1x digital and 4x zoom factor, respectively for maximum visualization of microglial process arbors, with 15-30 microglia present within each image. Sholl analysis was performed to quantify microglial arbor complexity within each region and between genotypes. Prior to the analysis, the z-stacks (approximately 30-50 Z planes) were projected into a maximum intensity image. A subset of microglia per image were selected, numbered, thresholded, and binarized using the moments thresholding method (Tsai) which most faithfully preserved overall microglial morphology. The Sholl analysis plugin (see Whitelaw et al. 2020, ImageJ) utilized concentric circles, drawn over each microglial arbors, radiating from the soma at 2μm intervals until the furthest concentric circle encompassed the whole microglia. The average number of intersections between the microglial arbor and each concentric circle, per image, was quantified, with approximately 15-30 microglia analyzed per image. The resulting Sholl curves for each microglia were individually integrated to calculate the average total number of microglial intersections per animal.

### Microglial density and distribution

Epifluorescent images were collected (microglia in GFP channel and nuclei in the DAPI channel) utilizing a Zeiss Axioplan 2 microscope (apochromat 10x/0.45 objective) and captured with an ORCA-spark digital camera (Hamamatsu), collected in Slidebook 6.0.24. Layers 1-6 of visual cortex as well as the ML and GCL in cerebellum were outlined in ImageJ using the polygon tool, and the area was measured. The cell counter tool was used to mark and number the position of each microglial soma. Results from 3-5 sections per animal were averaged. Density per region was calculated at the number of microglia per area in μm^2^. A nearest neighbor (NN) index for each cell was calculated by inputting soma positions into a custom MATLAB script ^72^. The spacing index, a density independent measure of microglial distribution was defined as NN^2^/density in each animal ^29^.

### Flow cytometry, and FACS

Flow cytometry and fluorescence activated cell sorting (FACS) were adapted from Karaahmet et. al. 2022. Mice were euthanized with an overdose of sodium pentobarbital and transcardially perfused with 0.1 M PBS. The brain was extracted, and the cerebellum and cerebral cortex were dissected and homogenized in 3mL ice cold FACS buffer (1X PBS + 0.5% BSA). The resulting homogenate was then filtered through a 70 μm cell strainer, which was then washed with a further 3mL ice cold FACS buffer. The suspension was centrifuged at 400 x g for 5 minutes at 4° C. The supernatant was discarded, and the remaining cell pellet suspended in a 40% Percoll PBS solution. These were then centrifuged again at 400xg at 4° C for 30 minutes with no break. The supernatant and resulting myelin were then aspirated. 70μL Fc Block (Biolegend) was then added, and the cells resuspended. Cells were then transferred to a 96 well plate for incubation with conjugated antibodies for flow cytometry and FACS, along with 100μL FACS buffer. Antibodies used were: CD11b-FITC (M1/70, Biolegend), CD45-APC/Cy7 (30F11, Biolegend), 7AAD (Invitrogen; cell death marker), P2Ry12-APC (S16007D, Biolegend), TMEM119-PE (106-6, Abcam, ab225496) and CD68 (137010, BioLegend). The plate was then centrifuged at 400xg for 5 min at 4° C, the supernatant discarded, the cells resuspended in 300μL FACS buffer and transferred to FACS tubes for sorting. Samples were sorted into Eppendorf tubes containing 300μL RLT buffer with ;-mercaptoethanol (BME, Sigma) (10μl BME/1mL RLT). Samples were kept on ice until sorting. Upon completion of collection samples were immediately frozen on dry ice. All males of both genotypes were sorted on the first day; all females of both genotypes were sorted on the second. Samples were kept at -80° C until submission for RNA sequencing.

### RNA-sequencing

Sorted cells were collected in 300 μL RLT Buffer (Qiagen) containing 2-mercaptoethanol (1µL/100µL RLT) and total RNA was isolated using the RNeasy Plus Micro Kit (Qiagen). RNA concentration was determined with the NanopDrop One spectrophotometer (NanoDrop) and RNA quality assessed with the Agilent Bioanalyzer 2100 (Agilent). 500 pg of total RNA was pre-amplified with the SMARTer Ultra Low Input kit v4 (Clontech) per manufacturer’s recommendations. The quantity and quality of the subsequent cDNA was determined using the Qubit Fluorometer 3.0 (Life Technologies) and the Agilent Bioanalyzer 2100 (Agilent). 150 pg cDNA was used to generate Illumina compatible sequencing libraries with the NexteraXT library preparation kit (Illumina) per manufacturer’s protocols. The amplified libraries were hybridized to the Illumina flow cell and sequenced using the NovaSeq6000 sequencer (Illumina) with target depth of 50 million with 50nt paired end reads per sample. Four samples, each representing individual animal, were sequenced per sex per treatment.

### Bioinformatics analysis

Raw reads generated from the Illumina basecalls were demultiplexed using bcl2fastq v19.1. Quality filtering and adapter removal was performed using FastP v.0.23.1. Processed reads were then mapped to the human reference genome (GRCm39 + gencode-M31 annotation) using STAR_2.7.9a. Read counts were quantified using both Subread-featureCounts v2.0.1 with “-s 0” indicating unstranded reads, and Salmon v1.5.2. All downstream analysis was carried out within R v4.2. Differential expression analysis was performed using DESeq2 with gene-level counts from featureCounts. Groups compared were cortex vs. cerebellum for each genotype and sex; and BL6 vs. P2RY12KO for each brain region and sex. Gene ontology analyses were performed using the clusterProfiler package. For all analyses, padj < 0.05 was use as a cut-off for statistical significance.

### Cranial window surgery

Cranial windows were created over both cerebellum and cortex adapted from the procedure described in Stowell et al 2018. Briefly, animals were anesthetized using a premixed cocktail of fentanyl (0.05 mg/kg), midazolam (5.0 mg/kg) and dexmedetomidine (0.5 mg per kg), in 0.9% saline administered intraperitoneally at 0.125mg/kg. Lubricant ointment was applied to the eyes and body temperature was maintained at 37°C. Aseptic technique was adhered to during all procedures: all tools were autoclaved for sterilization, a bead sterilizer was utilized between animals, and tools were kept in 70% ethanol during procedure. Mice were mounted in a stereotactic frame and head fixed for stability during surgery. The fur was removed, and the surgical area washed twice with 70% ethanol and betadine surgical scrub. An incision was made in the scalp and the skull cleared of membranes. In the case of cerebellar windows, a small amount of muscle was excised to expose the skull above cerebellum. Ferric sulfate, a veterinary vasocontractant (Kwik Stop), was used to staunch any bleeding that occurred, as was particularly frequent in P2Y12KO mice. A 3mm biopsy punch (Integra) was then used to create a circular score in the skull over either cortex or cerebellum. A 0.5mm drill bit was then used to create the window by tracing the score with minimal pressure. The skull was then removed. A glass cranial window composed of a 5mm coverslip and two 3mm coverslips (Warner instruments), attached with UV glue (Norland Optical Adhesive, Norland Inc) was then carefully lowered into the craniotomy. The window was secured and bonded to a headplate using C&B Metabond dental cement (Parkell Inc). The cement was used to cover the skull and seal the remaining surgical incision site. Slow-release buprenorphine was then administered subcutaneously as an analgesic, and the animals were monitored every 24hrs for 72hrs for signs of post-surgical pain.

### In vivo two-photon imaging

After a recovery period of at least 14 days to reduce inflammation, animals with cranial windows were prepared for in vivo two-photon imaging. A custom two-photon laser-scanning microscope was used (Ti:Sapphire, Mai-Tai, Spectra Physics; modified Fluoview confocal scan head, x20 objective lens, 0.95 NA, Olympus). Excitation for fluorescence imaging was achieved with 100-fs laser pulses (80 MHz) at 920 nm for GFP. All studies were carried out using anesthetized mice using the fentanyl cocktail described above. During imaging sessions and post-imaging recovery, mice were kept at 37 °C until they were alert. Imaging was conducted at 1x to 5x digital zoom with a 1-μm z step. Acute time-lapse imaging was carried out at 5-min intervals over 1h. Chronic imaging was carried out at day 1 and at day 7 at a 1x or 3x digital zoom. Image analysis was done offline using ImageJ and MATLAB with custom algorithms as described in Whitelaw et al. 2021.

### Process motility and surveillance

In vivo 50μm Z stacks were collected in the cerebellum and visual cortex in both P2Y12KO CX3CR1-GFP and CX3CR1-GFP every 5 minutes for 1hr producing 12 timepoints. Images were corrected for motion artifact in 3 dimensions and photobleaching offline in ImageJ with a custom macro. For analysis, 30μm max intensity z projections were made. As described in Whitelaw et al. 2020, for each time series image, a threshold was manually selected to include fine processes while excluding background noise. Binarized images of consecutive time points were then overlaid. Using a custom MATLAB script, consecutive time points were compared: pixels present in the first time point and absent in the second time point were defined as retractions; pixels absent in the first time point and present in the second time point were defined as extensions; pixels present in both were defined as stable. The retractions and extensions were summed and divided by the stable pixels for each of these consecutive timepoint pairs to generate motility indices for each of the pairs. The presented motility index represents the average of the motility indices calculated for each of the pairs within each animal. To calculate surveillance, or the total area surveyed by microglia across the imaging session, the binarized image was projected in the time dimension, and the fraction of pixels that were positive was determined ^15^.

### Cerebellar microglial soma motility

Z-stacks of 100μm of the same region were taken as described above on day 1 and day 7, using the vasculature to locate the same region. The positions of all visible microglial somal positions in the 100μm stack on each day were recorded in 3 dimensions, as were the positions of key features in the vasculature. A translational matrix was derived in MATLAB using the vasculature feature positions between days and applied to soma positions on day 7 to align the two timepoints. To obtain a quantification of microglial somal motility, a nearest neighbor quantification was utilized as described in Stowell et al. 2018. The coordinates of each microglial soma at day 1 were compared the transformed soma positions at day 7 in MATLAB and the nearest soma was found. The vector distance was then found between the 2 timepoints and recorded as microglial displacement. Outliers were removed as they often constituted the edge effects of subtle shifts in the image location between days. These displacements are presented either averaged by mouse or plotted on a frequency distribution. It is worth noting that this method underestimates the amount of somal movement as the NN distance is reflective of the nearest cell, which may not always represent the same cell across imaging days.

### Laser ablation injury and analysis

Laser ablation injuries were induced by tuning the laser to 800 nm, increasing the laser power to 5W, and performing a point scan on an unlabeled area in the parenchyma for 2-15 s. This generated an auto-fluorescent injury core indicative of tissue damage. After induction of the ablation injury, 60μm z-stacks were taken every 5 min for 1hr. Analysis of these images was performed in a very similar manner to that described in Whitelaw et a. 2021. In brief, two aspects of the microglial chemotactic response were analyzed: the convergence of microglial processes on a central point, in this case an injury or compound point source, and process movement toward that source. Maximum z projections were created of each microglial ablation response from the +/-10μm around the injury core. A polygon was outlined around the autofluorescent core generated by the laser injury, and this area was excluded from analysis. A donut shaped region of interest (ROI) with a of thickness 40 pixels (10μm) was generated around the injury core from that outline polygon. Convergence was calculated as the fraction of this ROI that was occupied by positive pixels, subtracted by the initial occupancy. To make statistical comparisons across groups, convergences at specific timepoints were compared, denoted in relevant figures. To better capture microglial process movement throughout the entire field of view, optic flow-based approach was utilized. This approach compares two consecutive frames and determines, for each pixel, the direction and magnitude of motion in the image. To find the degree to which this movement was directed toward the site of injury, the velocity vectors were projected onto the normalized vector pointing toward the injury, further generating a measurement of directional process velocity. In order to isolate microglial movement, a binary mask based on the thresholded images was applied to the vectors. Finally, the average directional velocity was calculated across all microglial pixels. In order to compare across groups, process velocity was averaged 10 minutes after injury or puff application until the response was complete (40-50 minutes *in vivo*, 20 minutes in slices). This excluded initial artifacts caused by tissue deformation as well as the period after convergence has largely completed. To better describe microglia process behavior at a greater distance from the ablation core, an additional analysis was performed as described in Cealie et. al. 2024. The polygon tool in ImageJ was used to an ROI around the core and then around the ring created by the microglial processes as they approached the injury core, with the polygon going back to the image border when gaps in the ring occurred, especially at earlier time points. Areas were then measured in microns for all ROIs and the measurements saved. The core area was subtracted at each time point and then that value was normalized by dividing that value the initial area of the process ring at t=0.

### Acute slice preparation

For all slice experiments, slices were imaged between 1-4h from the time of slice preparation to avoid overt microglial reactivity. For focal stimulation experiments in acute slices, mice were decapitated, and the brain was rapidly removed into ice-cold HEPES-buffered ACSF (artificial cerebrospinal fluid), containing the following: 140 mm NaCl, 5 mm KCl, 1 mm NaH2PO4, 10 mm glucose, 2 mm MgCl2, 10 mm HEPES, and 2 mm CaCl2; pH adjusted to 7.4 with 5 M NaOH; and oxygenated with 100% O2 (HEPES-ACSF; ∼297 mOsm). Horizontal sections of 400μm thickness encompassing cerebellum or coronal sections of 400μm thickness encompassing visual cortex were prepared using a vibratome (Vibratome 1000), with the brain submerged in ice-cold ACSF. Slices were transferred to a recovery chamber containing HEPES-ACSF at 33°C. HEPES-ACSF (heated to 34–36°C) was also used as the extracellular solution during imaging. For imaging, slices were placed in a perfusion chamber (RC-27L, Warner Instruments), continuously perfused with ACSF at a rate of ∼2 ml/min and imaged at a depth of ∼50– 100 μm to avoid areas damaged from the slicing procedure ^15,40^.

### Focal ATP release

A 1 μM ATP (Sigma), 4% tetramethylrhodamine (“rhodamine”; average molecular weight 4400; Sigma) solution was prepared in HEPES-ACSF and backfilled into a glass microelectrode (∼3–4 MΩ resistance; BF150-110-10 glass pulled with P-97 puller, Sutter). The glass microelectrode was slowly inserted into the tissue using a micromanipulator (MP225, Sutter). The end of the pipet was connected to a Picospritzer III (Parker). To image the microglial response around the pipette tip, an initial z-stack was obtained, then a pressure pulse (100 ms, 15 psi) was released ATP onto the slice during the second time frame (t = 0 min). Additional z-stacks were acquired for 30 min (every 1 or 2 min). In addition to ATP, solutions of rhodamine dye and either potassium chloride (140 mM, Sigma), sodium glutamate (100 mM, Sigma) and purified HMGB1 (10nM, Sigma) were all prepared in ASCF and applied in the same manner.

### Analysis of directed motility toward a point source

To analyze the microglial response to focal stimulation, the protocol used for injury analysis was modified, in a similar manner to that described in Whitelaw 2020. In brief, a point was placed at the end of the pipet tip, rather than a polygon around the injury core. Next, a polygon was drawn around the pipet, to exclude it from the analysis. For convergence, a circular ROI of radius 50 pixels (15μm) centered on the tip of the pipet was used as the area in which to calculate microglial occupancy, instead of the donut ROI around the injury core. Over time, microglial processes often entered the pipet tip, thus leading to a decreased value of convergence. Thus, to compare across groups, the maximum convergence was used in the pipette experiments.

### Delay eyeblink conditioning

Delay eyeblink conditioning (dEBC) is a well characterized and studied paradigm of cerebellar learning. During the task, a neutral, conditioned stimulus (CS, an LED at 405nm) and a noxious unconditioned stimulus (US, and air puff at 20 PSI to the cornea) are paired, while mice are allowed to run freely on a wheel. The experimental set up used here is similar to what is described in Broussard et al. 2022, but without the ability to monitor motor activity. Male mice of either P2Y12KO experimental or BL6 control genotype were dark reversed 14 days before experiment start to align mouse circadian rhythms and activity patterns with the experiment ^43^. To implant headposts, mice were anesthetized with the fentanyl cocktail described above and prepared for surgery as described above for cranial window implantation. Once the skull was exposed, the membranes over the skull were cleared using Kwik Stop styptic gel and 70% ethanol solution. Headposts were secured to the skull using C&B Metabond dental cement (Parkell Inc). The cement was used to cover the skull and seal the remaining surgical incision site. Slow-release buprenorphine was then administered subcutaneously as an analgesic, and the animals monitored every 24hrs for 72hrs for signs of post-surgical pain. After a 5-day recovery period, mice were trained in wheel running and head fixation for 5 days. Training occurred in low light conditions on running wheels identical to the experimental. Training consisted of sessions of head fixation and wheel running of increasing duration on each training day (15 minutes days 1 and 2, 20 minutes day 3, 25 minutes day 4, 30 minutes day 5).

The experiment began the day after the fifth day of training and continued for 10 days. During this time mice were head fixed and allowed to run freely on a running wheel. Each day, mice were presented with a blue light (conditioned stimulus (CS)) followed after a short delay by an air puff to the eye (unconditioned stimulus (US)), and the blink response was recorded. Each trial was 1000ms long. A typical trial consisted of a 100ms pre-CS period, a 280ms CS period, the last of which overlapped with 30ms in which the US was presented. 220 trials were recorded per day with 10% of trials using the CS alone. A random inter trial interval of 5000ms-12000ms was used so as not to habituate the mice to stimulus timing. Eye recordings were made in the dark with the eye illuminated with an infrared lamp. Video oculography was used to capture eye movements during the trail. Recordings and stimuli were created using a Raspberry pi and Arduino microcontroller respectively.

Eye recordings were binarized with a semi-automated thresholding algorithm using a python script provided by Dr. Joey Broussard ^46^. The animal eye region of interest was identified at trial outset by the experimenter and was then automatically thresholded. Changes in pixel value caused by eyelid closure during the blink response were recorded as eye position. This was done for all animals on all experimental days. If eyelid closure achieved a specified threshold (25% of the full blink response) during the conditioned response (CR) window of 150ms-350ms then the trial was counted as a CR. The percent CR rate was recorded on each day as total number of trials in which CR was achieved divided by the number of trials. A custom MATLAB script was used to generate the percent CR.

### Statistical analysis and notes

Appropriate statistical methods were used to make comparisons between groups and reported as such. An α of p<0.05 was treated as significant. All analyses and drug treatments were performed blinded to genotype and condition where applicable. Statistical analyses were performed in GraphPad Prism version 10.0.0 for Mac (GraphPad Software, Boston, Massachusetts USA, www.graphpad.com). The code/software used in the analyses presented here are freely available online at https://github.com/majewska-lab.

## Supporting information

Supplemental Movie 1 CTX CTRL

Supplemental Movie 2 CBL CTRL

Supplemental Movie 3 CTX P2Y12KO

Supplemental Movie 4 CBL P2Y12KO

Supplemental Movie 5 Velocity Vectors

## Acknowledgements

We thank Dr. Joey Broussard for his assistance and generosity in the set of the dEBC rig and in the analysis. We thank Dr. Kaye Thomas, Dr. Julie Zhang of the University of Rochester Confocal light Imaging core and the University of Rochester the flow cytometry core facility. We would like to thank Dr. Farrell “Ric” Robinson for his guidance in the early stages of this study. We would thank Cassandra Lamantia for her assistance and maintenance of many of the animal strains used here. This work was supported by F31NS120609 (MBS), R01EY019277 (A.K.M), R21NS099973 (AKM), R01AA02711 (AKM), NSF000497 (AKM).

## Contributions

MBS and AKM planned and designed the experiments. MBS carried out experiments, analyzed data. MBS wrote the manuscript with editorial assistance from AKM. RDS provided training, feedback in designing experiments and provided data in Figure 8. RLL helped devise, carry out, and analyzed the data from the behavioral experiments in Figure 10. LL helped plan and perform microglial isolation, RNA sequencing, and subsequent analysis appearing in Figures 2-6. ANV collected and analyzed data presented in Figure 1. BSW assisted in training and devised the analysis used in Figures 8 and 9. BSW further provided data appearing in Figure 8.

---

**Supplemental Figure 1.**
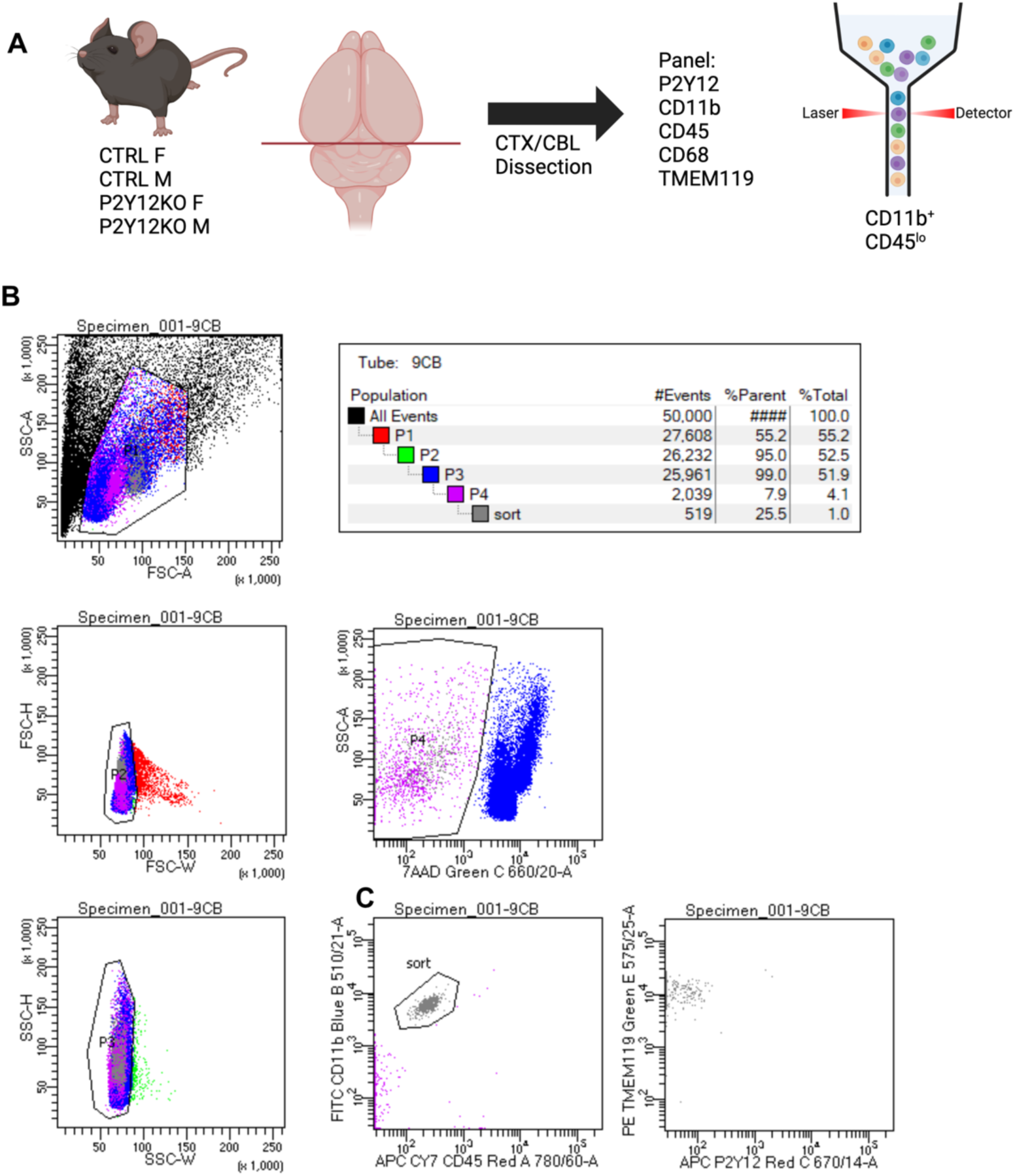
Experimental design and gating strategy used in in microglial isolation via FACS. Schematic illustrating experimental design (A). Brain Tissue was harvested from male (M) and female (F) P2Y12 deficient (P2Y12KO) and BL6 control (CTRL) mice. Cerebral cortex (CTX) and cerebellum (CBL) were dissected from each brain and homogenized. Myelin was removed using a Percoll gradient. Cells were stained for a flow cytometry panel of microglial markers. Microglia were isolated using FACS, sorting for CD11b+/CD45lo (A). The 8 final experimental groups were as follows: CTRL CTX F (n=4), CTRL CBL F (n=4), P2Y12KO CTX F (n=4), P2Y12KO CBL F (n=4), CTRL CTX M (n=4), CTRL CBL M (n=3), P2Y12KO CTX M (n=4), P2Y12KO. Gating strategy employed when sorting microglia sorting for CD11b+/CD45lo (B and C). Panel A created in Biorender.

**Supplemental Figure 2.**
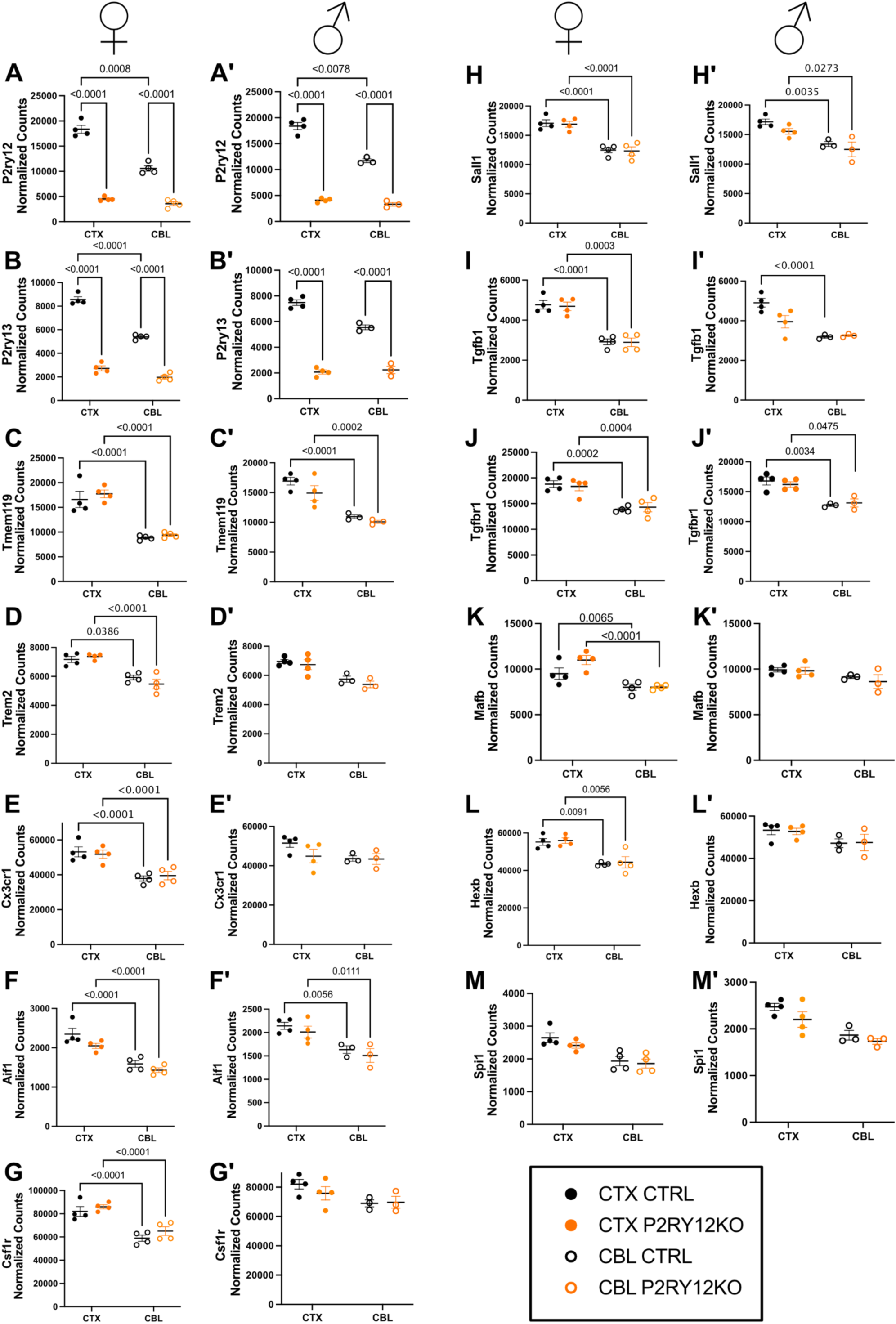
Individual microglial signature gene transcript levels are modulated regionally, with minor effects of P2Y12 deficiency. Microglia were sorted from cortex (CTX n=4, closed circles) and cerebellum (CBL n=3, open circles) from control (CTRL, black) and P2Y12 deficient mice (P2Y12KO, orange), from female and male mice. Prime (‘) denotes males. All data were compared using padj <0.05 for statistical significance.

**Supplemental Figure 3.**
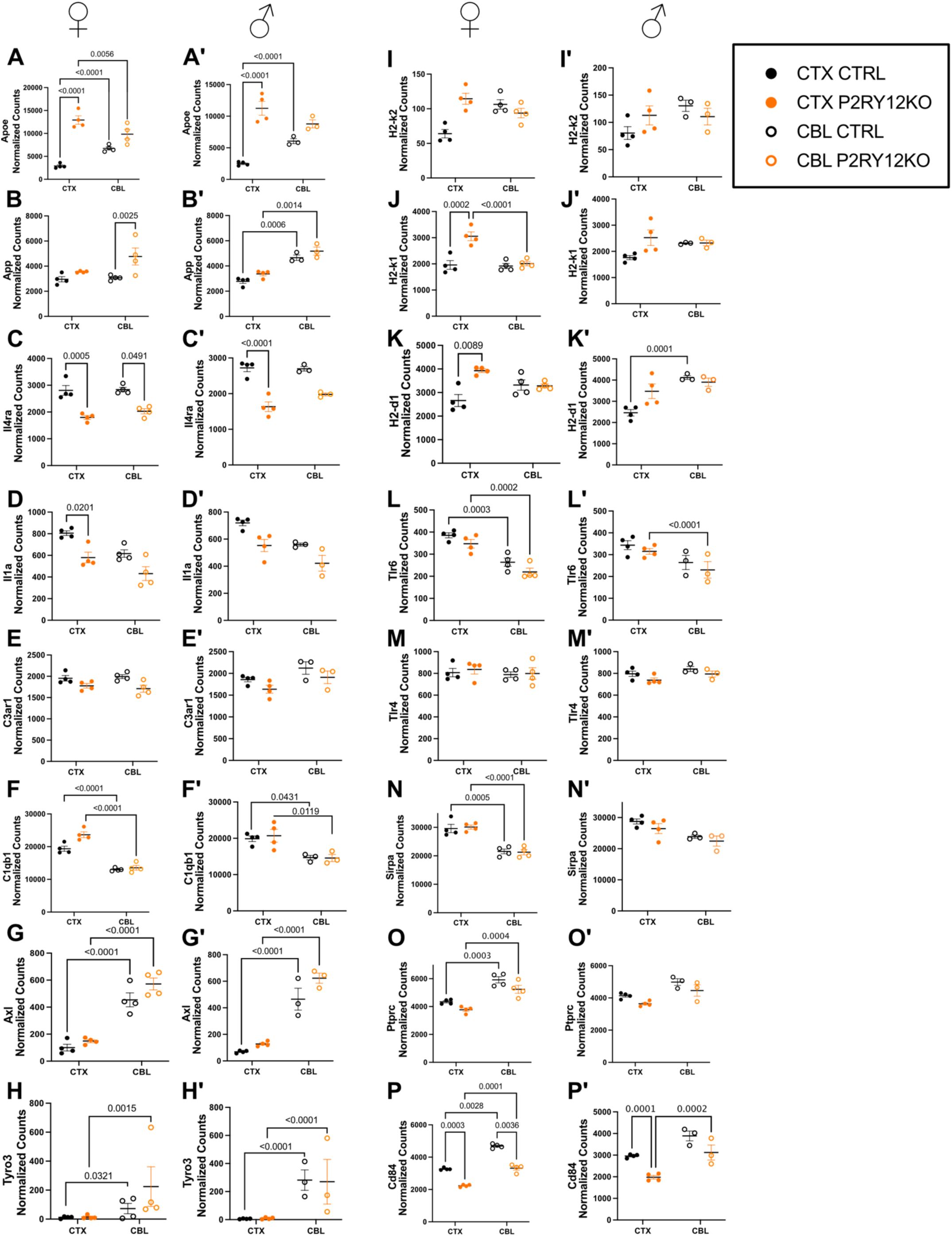
Key inflammation signaling gene transcript levels are modulated by both region and P2Y12 deficiency. Microglia were sorted from cortex (CTX n=4, closed circles) and cerebellum (CBL n=3, open circles) from control (CTRL, black) and P2Y12 deficient mice (P2Y12KO, orange), from female and male mice. Prime (‘) denotes males. All data were compared using padj <0.05 for statistical significance.

**Supplemental Figure 4.**
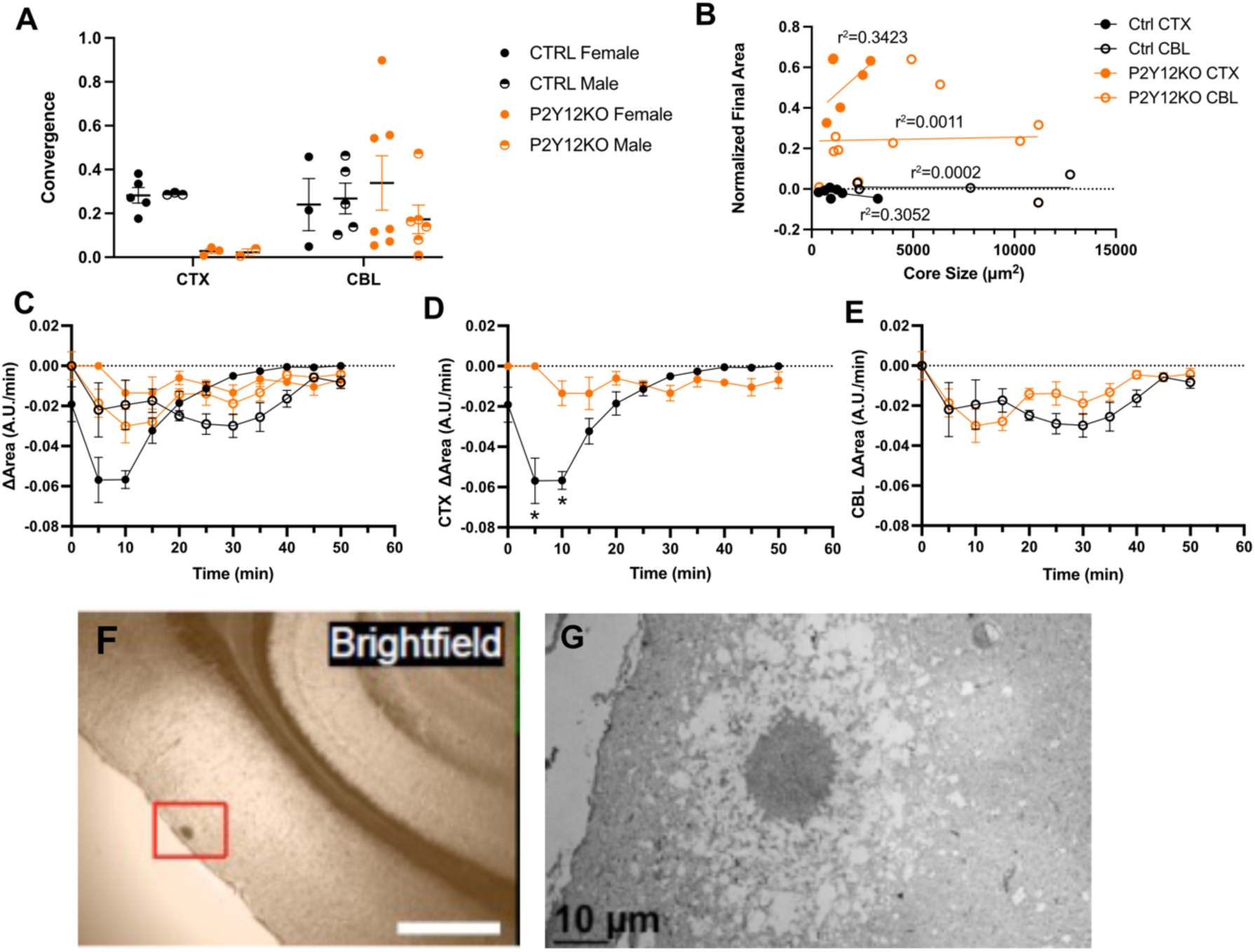
Microglial injury response dynamics in the absence of P2Y12 in the cortex and cerebellum. The microglial injury showed no effect of genetic sex on final convergence (three-way mixed measures ANOVA genotype p=0.0996, region and genotype p=0.0932, Bonferroni post-hoc) (A). Normalized final area as a function of initial core size; the ablation response occurred independent of core size, except for a positive correlation in the P2Y12 cortical (CTX) group (simple linear regression) (B). Rate of change in normalized area (ΔArea) as a function of time in all groups (C). P2Y12KO CTX differed significantly from CTRL CTX in early ablation response (two-way ANOVA, significant effect of genotype, p<0.0001 and time p<0.0001; genotype and time p<0.0001; Bonferroni post hoc * p<0.00001) (D). CBL P2Y12KO response displayed slightly altered ΔArea over a time (two-way ANOVA, significant effect of genotype, p=0.0421 and time p=0.0010; Bonferroni post-hoc) (E). Brightfield image of a CTRL CTX ablation in fixed tissue, area in inset was prepared for imaging and reconstruction with serial electron microscopy (EM) (F). EM image showing extensive damage around ablation core (G scale bar = 10μm).

